# Plasmids promote antimicrobial resistance through Insertion Sequence-mediated gene inactivation

**DOI:** 10.1101/2025.08.12.669853

**Authors:** Jorge Sastre-Dominguez, Paloma Rodera-Fernandez, Javier DelaFuente, Sandra Martínez-González, Susana Quesada, Marina Valencoso-Requena, Alicia Calvo-Villamañan, Coloma Costas, Ayari Fuentes-Hernández, Alfonso Santos-Lopez, Alvaro San Millan

## Abstract

Antimicrobial Resistance (AMR) is a major threat to public health. Plasmids are mobile genetic elements that can rapidly spread across bacterial populations, promoting the dissemination of AMR genes in clinical bacteria. In addition, plasmids are enriched in insertion sequences (IS), which are small transposable elements able to translocate between genetic locations. Importantly, IS transpositions commonly lead to gene inactivation, which can in turn promote AMR (e.g. through the modification of the antibiotic target). In this study, we combined experimental, bioinformatic and computational approaches to investigate the role of plasmids as catalysts of AMR through IS-mediated gene inactivation. Our results revealed that plasmid pOXA-48, which encodes two IS1 elements, increases the rate of resistance acquisition to multiple antibiotics in clinical strains of *Klebsiella pneumoniae* through IS1-mediated gene disruption. Moreover, a large screen of genome databases confirmed that the inactivation of genes through plasmid-encoded IS elements is an extended mechanism of AMR evolution. Finally, both our experiments and computational model revealed that conjugative plasmids can promote this route of AMR acquisition while invading complex bacterial communities. Overall, our study reveals that conjugative plasmids fuel AMR not only through the dissemination of resistance genes, but also through IS-mediated gene inactivation, promoting the evolution of multidrug resistance in bacteria.

## Introduction

Antimicrobial resistant bacteria represent one of the most critical public health menaces^1,2^, threatening to escape the current available treatments against bacterial infections^3^. The remarkable success of bacteria against antibiotics can be explained by the wide range of molecular mechanisms through which they rapidly evolve antimicrobial resistance (AMR)^4,5^. These mechanisms are acquired via two fundamentally different evolutionary routes: mutations of pre-existing genes or the acquisition of novel DNA through horizontal gene transfer (HGT)^6–8^. In this context, mobile genetic elements (MGE) have received major attention due to their key role as vehicles for the dissemination of AMR genes through HGT^9^.

Conjugative plasmids stand out as the main spreaders of AMR in clinical settings^7,10^. The dissemination of AMR mechanisms by conjugative plasmids has played a crucial role in the emergency of high-risk nosocomial pathogens^10–12^. Moreover, recent work has demonstrated that plasmids can also foster bacterial evolution beyond HGT^13,14^. For example, plasmids can alter the transcriptional profiles of their hosts^15–18^, promote genetic variability^19,20^ and even increase the mutation rate of plasmid^21^ or chromosomal genes^22^. This is especially worrisome in the clinical context, where multidrug resistant (MDR) bacteria can arise from particular plasmid-bacteria associations which become extremely successful^23,24^. A good example is the association between carbapenem-resistance plasmids and certain Enterobacterales clones (e.g. pOXA-48 and *Klebsiella pneumoniae* ST11^25^).

Other MGEs, such as Insertion Sequences (IS), also play a key role as vehicles of AMR in bacteria^26,27^. IS are transposable elements that encode the information necessary for their own transposition^28^. IS are also able to transpose resistance genes, which they can mobilize between plasmids, or between plasmids and chromosomes^29,30^. However, as conjugative plasmids, IS also play a role in the evolution of resistance beyond gene transfer^31^. For example, the insertion of IS elements can modify the expression of nearby genes or inactivate them by disrupting their coding sequence, which can affect bacterial metabolism, virulence and resistance^31–33^. Crucially, gene inactivation is a common mechanism leading to AMR in clinically relevant bacteria. For example, gene disruption can lead to resistance phenotypes through efflux pump overexpression^34^, reduced cell permeability^35^, or drug target modification^36^. One particularly alarming example of this phenomenon is the rise of IS-mediated colistin resistance (acquired predominantly by the disruption of the *mgrB* gene^37–39)^ in carbapenem-resistant Enterobacterales (CRE)^40,41^. This combined phenotype impacts the available therapeutic options dramatically, as it limits the efficacy of two last-resort antibiotics of critical importance.

Plasmids are enriched in IS elements^28,42^. Consequently, plasmids can provide benefits to their bacterial hosts not only by disseminating beneficial genes, but also by increasing the mutation rate of their hosts by IS transposition^43^. Recent results from our lab with a local collection of clinical Enterobacterales showed that the worldwide distributed AMR plasmid pOXA-48^44^, not only confers carbapenem resistance, but also allows for the rapid pathoadaptation of clinical enterobacteria through the transposition of plasmid-encoded IS1 elements^45^. In light of all this evidence, we hypothesize that plasmids could promote AMR not only through AMR gene dissemination but also through chromosomal gene inactivation. To experimentally test our hypothesis, we used a model system based on the plasmid pOXA-48 and multiple clinical Enterobacterales. We combined several experimental approaches to demonstrate that pOXA-48 accelerates the rate of AMR evolution to a wide range of antibiotics through IS1-mediated chromosomal gene disruption. Crucially, to expand our results beyond the experimental system, we analyzed more than 50,000 bacterial genomes from public databases, which allowed us to link the presence of plasmid-encoded IS with the acquisition of AMR through IS-mediated chromosomal gene inactivation across multiple species. And finally, we used computer simulations to explore the general impact of plasmid conjugation and plasmid-borne IS transposition on the evolution of AMR in bacterial populations. Overall, our results demonstrate that plasmid-mediated IS gene inactivation promotes the acquisition of AMR in bacteria.

## Results

### pOXA-48 promotes the acquisition of colistin resistance

To analyze the effect of plasmid-encoded IS elements on the evolution of AMR through chromosomal gene inactivation, we used the globally distributed carbapenem resistance plasmid pOXA-48. pOXA-48 only confers carbapenem resistance and encodes two copies of the IS1 element (**Fig. 1A**). First, we generated pOXA-48-carrying and pOXA-48-free isogenic pairs in 5 clinical strains of *K. pneumoniae*. These strains were selected upon their clinical relevance, phylogenetic diversity (ST15, ST377, ST432 and ST1427; **S. Table 1**) and susceptibility profiles to multiple antibiotics (see Methods). Then, we set up a fluctuation assay^46^ to test whether pOXA-48 carriage led to increased colistin (COL) resistance. Briefly, we cultured independent replicates of each strain (n = 35) in the absence of antibiotics, allowing for random mutations or IS transpositions to arise in absence of selection. We then plated the cultures with and without COL pressure and calculated the phenotypic resistance mutation rate to COL^47^ (**Fig. 1B, S**. **Table 2**). As a control, we also tested the phenotypic resistant mutation rate to rifampicin (RIF), as resistance to this antibiotic cannot be achieved through gene inactivation^48^ (**S. Fig. 1**). We observed a significant increase in the COL resistance mutation rate in all the pOXA-48 carrying strains tested (**Fig. 1B**; p < 0.05, maximum likelihood ratio tests after Bonferroni-Holm adjustment), but no increase in the RIF resistance mutation rates (**S. Fig. 1**; p > 0.05, maximum likelihood ratio tests after Bonferroni-Holm adjustment). To confirm the role of IS1 in the increase in COL mutation rate, we built an IS1-less version of pOXA-48 (pOXA-48ΔΔIS1, see Methods, **Fig. 1B**). We introduced it in one of the clinical *K. pneumoniae* strains (KPN08) and repeated the fluctuation assays as described above (**Fig. 1B**). Interestingly, we observed no significant differences between the mutation rates of pOXA-48-free and pOXA-48ΔΔIS1-carrying strains in any of the antibiotics (p < 0.05, maximum likelihood ratio tests after Bonferroni-Holm adjustment).

**Figure 1.**
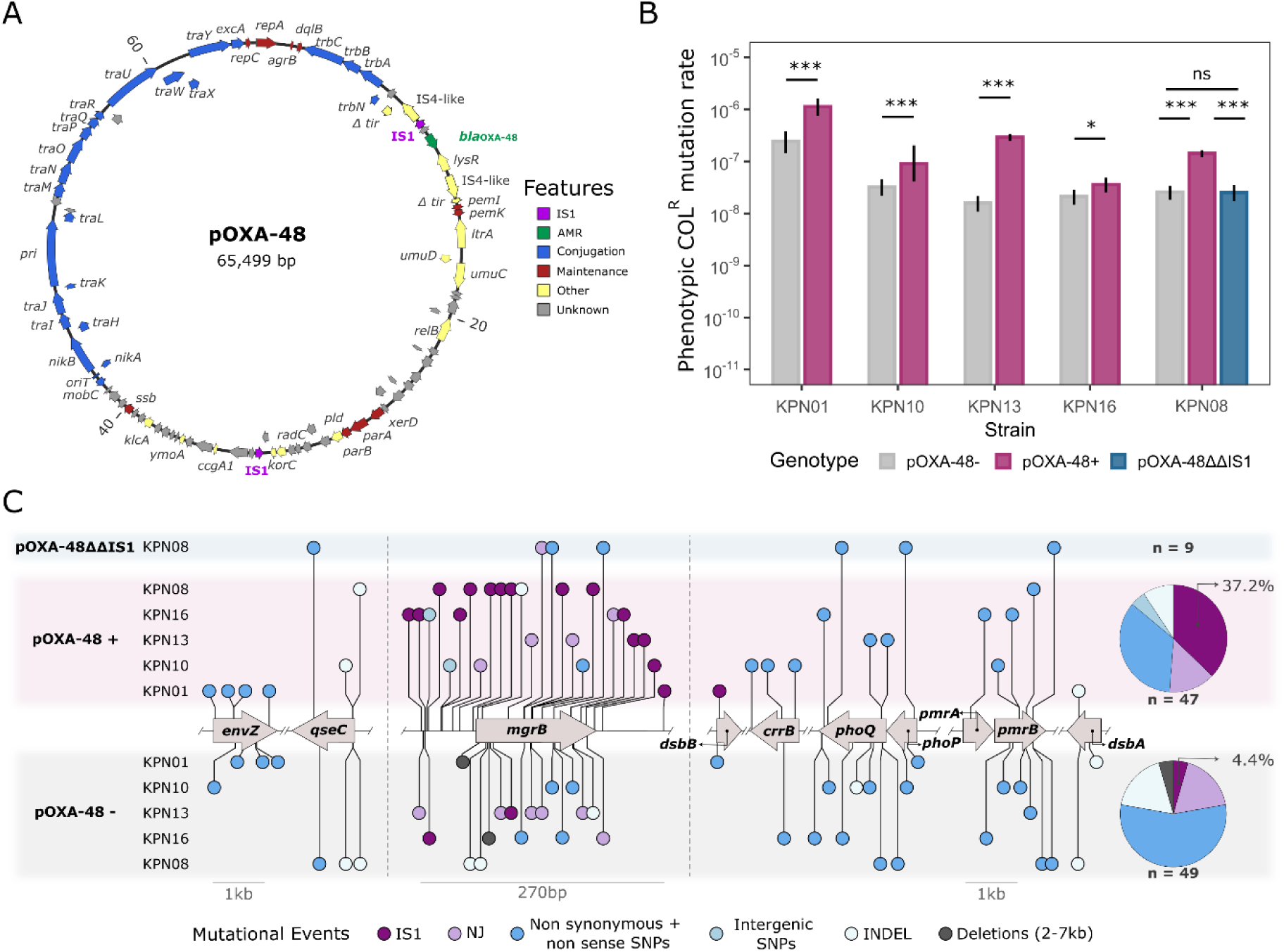
Resistance mutation rate to colistin associated with pOXA-48 carriage. **A)** Genetic map of the pOXA-48 plasmid used in this work (K8 variant^50^). Arrows represent the coding sequences of the genes (CDS), and the colors represent the function of each CDS. IS1 elements are highlighted in purple, and the carbapenem-resistance gene *bla*OXA-48 in green. Only the name of CDS with known function are shown. Genomic coordinates are represented in kilobase pairs (Kbp). **B)** Phenotypic resistance mutation rate to colistin (mutations per cell division) of the isogenic pairs of *K. pneumoniae* clinical strains either not carrying pOXA-48 (grey), carrying pOXA-48 (purple), or carrying the IS1-depleted version of pOXA-48 (pOXA-48ΔΔIS1; blue). The black bars indicate the 95% confidence interval. Asterisks denote statistical significance in the mutation rate between pOXA-48- and pOXA-48+ (and pOXA-48ΔΔIS1) for each pair based on maximum likelihood ratio tests (n = 35) after Bonferroni-Holm adjustment for multiple testing (p-value: 0 < *** < 0.001 < ** < 0.01 < * < 0.05 < ns). **C)** Mutations detected in the colistin resistant mutants. Mutations accumulated in the absence of pOXA-48 are represented on the lower part of the panel (grey background). Mutations accumulated in the presence of pOXA-48 (purple background) or pOXA-48ΔΔIS1 (blue background), are represented at the top of the panel. Pie charts represent the different types of mutations accumulated, as indicated in the legend (including those affecting non-represented genes). The number of strains sequenced for each condition is summarized in the right side of the panel. New junctions mediated by the IS1 are represented in dark purple while new junctions not mediated by the IS1 are in light purple. Dark blue indicates protein-altering SNPs, light blue represents intergenic SNPs, white denotes short INDELs, and gray corresponds to deletions affecting multiple genes (ranging from 2 kb to 7 kb). All genes of interest are plotted in order according to the reference genome, maintaining their relative positions while omitting the genetic regions between them. The scale is preserved for all genes except for *mgrB*, which is zoomed in to accurately visualize all events affecting the 144 bp long gene.

To analyse the molecular mechanisms underlying the COL resistance, we sequenced the whole genome of 105 COL resistant colonies from the different experimental treatments (one colony per independent replicate [n=7-10], per strain and version; **Fig. 1C**). Most sequenced clones differed from the susceptible parental genome only by a single mutation. The majority of mutations were found in 10 different genes previously associated with COL resistance, most of them related to the lipid A, which is part of the outer membrane lipopolysaccharide (LPS) and the molecular target of COL (**S. Table 3**). Crucially, we observed a significantly higher proportion of mutants with insertional inactivation caused by IS1 elements in pOXA-48-carrying strains (37.2%) than in pOXA-48-free strains (4.4%; p < 0.05, Fisher’s exact test after Bonferroni-Holm adjustment; **Fig. 1C; S. Table 4**). Interestingly, most insertions affected *mgrB* (84.8%), the negative regulator of the *phoPQ* two-component system, which controls the modification of lipid A^49^. We speculated that this bias in the insertion target could be caused by the IS1 predilection for AT-rich regions^28^, as *mgrB* was the target gene with the lowest GC content (∼40%; **S. Fig. 2**).

**Figure 2.**
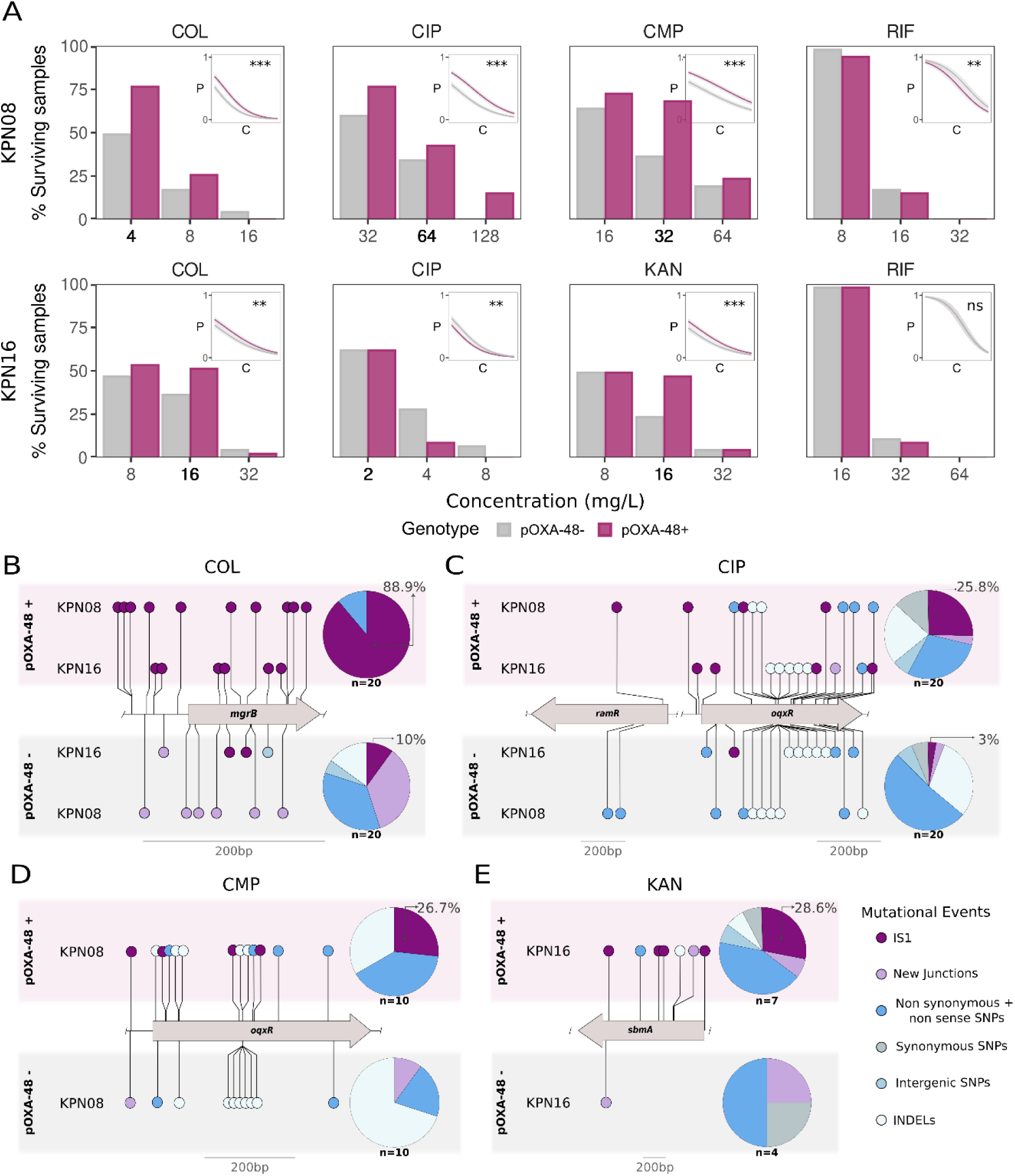
pOXA-48 promotes acquisition of resistance to multiple antibiotics. **A)** Surviving populations (%) in the assays (n = 47) at different antibiotic concentrations (in mg/L). Bar plot y-axis indicates the surviving replicates either carrying (pOXA-48+, purple) or not carrying pOXA-48 (pOXA-48-, grey) at different concentrations of each of the antibiotics (x-axis) for each of the *K. pneumoniae* strains tested (top: KPN08; bottom: KPN16). Darker bold concentrations in x-axis are those from which populations were sequenced. Inset plots show the predicted survival probability as a function of the increasing antibiotic concentration of colistin (COL), ciprofloxacin (CIP), chloramphenicol (CMP), kanamycin (KAN) and rifampicin (RIF). We predicted the probabilities by building a binomial logistic regression model per strain and antibiotic based on the number of surviving samples during the assays (n = 47). Asterisks denote statistically significant differences in the survival probability between both genotypic versions of the strains to each antibiotic (p-value: 0 < *** < 0.001 < ** < 0.01 < * < 0.05 < ns). **B-E)** Mutations detected in the common highest concentration for both conditions (pOXA-48 carrying and pOXA-48 free strains) where at least 10 replicates survived in the experimental assays for Colistin (**B**), Ciprofloxacin (**C**), Chloramphenicol (**D**) and Kanamycin (**E**). Mutations accumulated in absence of pOXA-48 are represented on the lower part of the panel (grey background) while those accumulated in presence of pOXA-48 are represented at the top of the panel (purple background). Only the genes showing parallel evolution among the samples and a disruption caused by an IS1 element are shown -full list of mutations are shown in **S.** Fig. 3-. Pie charts of all the events are displayed on the right for each condition (including those affecting non-represented genes). The number of strains sequenced for each condition is summarized in the right side of each panel, under each pie chart. New junctions mediated by IS1 elements are represented in dark purple while new junctions mediated by other elements are in light purple. Dark blue indicates protein-altering SNPs, light blue denotes intergenic SNPs, light grey represents synonymous SNPs, and white denotes INDELs. All genes of interest are plotted in order according to the reference genome, maintaining their reference positions while omitting the genetic regions between them.

### pOXA-48 enhances acquisition of resistance to multiple antibiotics

Gene inactivation can lead to different AMR phenotypes^26,27^. Therefore, we decided to expand our screen and study the effect of pOXA-48 on the acquisition of ciprofloxacin (CIP), chloramphenicol (CMP), and kanamycin (KAN) resistance. CIP, CMP and KAN belong to three different families of antibiotics of clinical relevance (fluoroquinolones, amphenicols, and aminoglycosides, respectively), and resistance to all of these antibiotics can be achieved through gene inactivation^34,35^. Fluctuation tests are low-throughput and laborious, so we decided to follow a different experimental approach. We exposed several independent colonies (n = 47) of two *K. pneumoniae* strains (KPN08 and KPN16), either carrying or not carrying pOXA-48, to multiple concentrations of each antibiotic (0.5x minimal inhibitory concentration [MIC], 1xMIC, 2xMIC 4xMIC). Again, we included RIF as a control. We selected KPN08 and KPN16 since they are susceptible to multiple of these antibiotics (KN08 to COL, CIP, CMP and RIF, and KPN16 to COL, CIP, KAN and RIF). After an overnight culture, we quantified the number of surviving populations in each antibiotic concentration (see Methods). This approach allowed a high-throughput exploration of resistance acquisition at different concentrations of several antibiotics. As predicted, we observed a larger number of surviving populations carrying pOXA-48 in comparison to their pOXA-48-free pair in most of the strain-antibiotic combinations (5/6), except for RIF (**Fig. 2A**).

To analyze the molecular mechanisms driving AMR, we sequenced the whole genome of 111 surviving populations from the different experimental treatments (COL, CIP, CMP and KAN, **Fig. 2B-E**). We selected, for each strain-antibiotic combination, the populations from the highest antibiotic concentration in which at least 10 replicates survived. Results showed a higher number of IS1-mediated gene inactivations in presence of pOXA-48 in the four antibiotics (**Fig. 2B-E, S. Tables 3 and 4**; p < 0.05, Fisher’s exact test after Bonferroni-Holm adjustment). Again, COL resistance was achieved by disrupting genes involved in the lipid A biosynthesis pathway (**Fig 2B**). CIP and CMP resistance were mostly acquired by the inactivation of RND efflux pump negative-regulators -*oqxR* and *ramR* for CIP and *oqxR* for CMP (**Fig. 2C-D**). KAN resistance was primarily achieved by the inactivation of the membrane transporter *sbmA* (**Fig. 2E**), and mutations on ABC transport systems (**S. Fig. 3**). These observations confirmed that pOXA-48 generally promotes the emergence of IS1-mediated AMR in *K. pneumoniae*.

**Figure 3.**
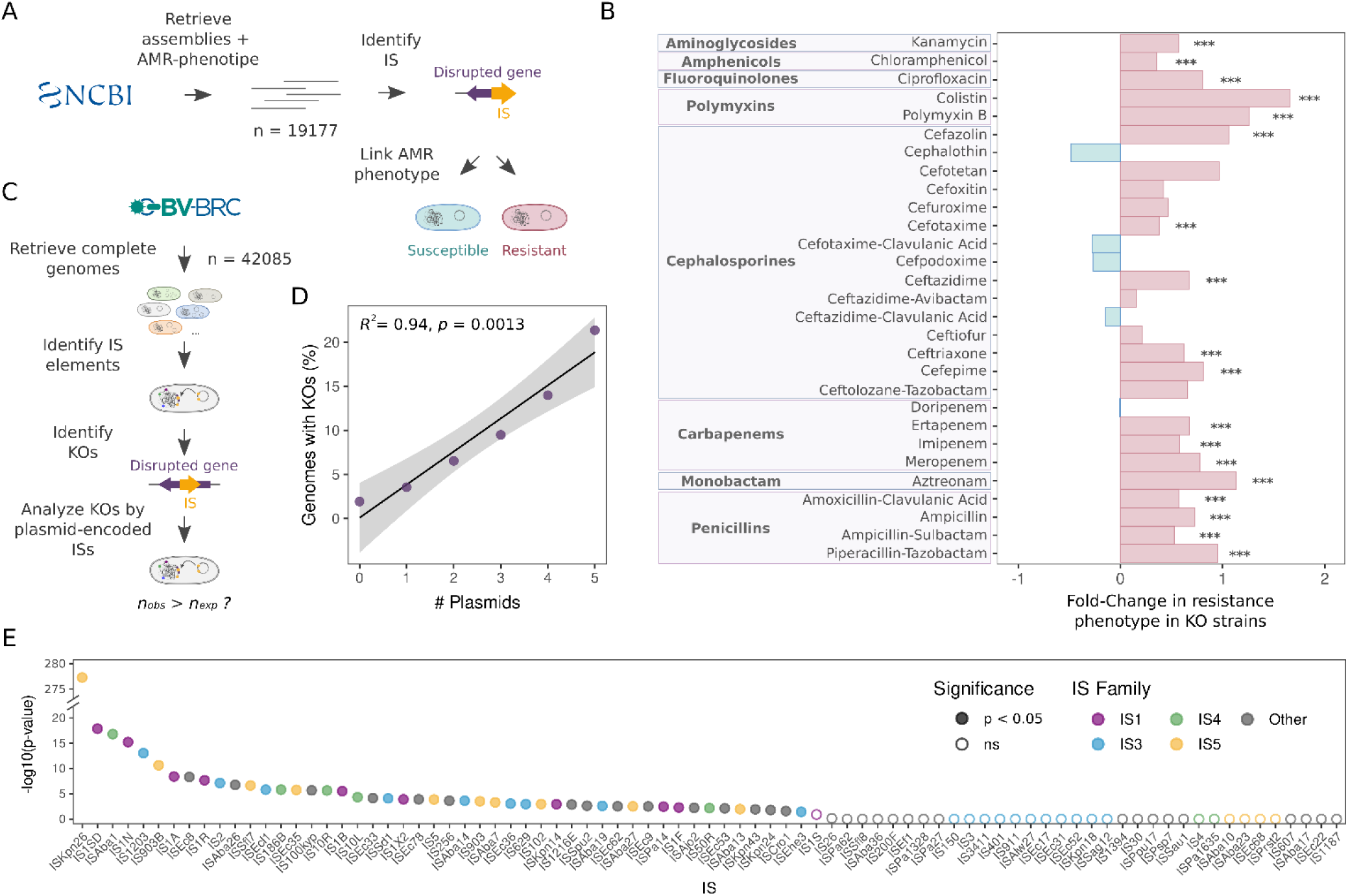
Plasmid-encoded IS elements promote AMR through gene inactivation in Enterobacterales. **A)** Scheme of the pipeline followed to analyze the relationship between IS-KO-AMR genes (experimental targets, genes affecting membrane composition and permeability, RND and MFS efflux pump genes and RND and MFS efflux pump regulators) and the AMR-phenotype of strains retrieved from the NCBI Pathogen Detection Database. **B)** Association between IS elements causing the disruption of KO-AMR genes and the resistance phenotype of the strain. X-axis indicates the Fold-Change (Log10 scale) in the frequency of strains showing resistant phenotype to each antibiotic when carrying IS-KO-AMR genes (relative to the strains not carrying KOs in KO-AMR genes). Asterisks indicate significant increase in the proportion of resistant strains associated with IS-KO-AMR genes. Red bars indicate that the proportion of AMR strains is higher for the strains showing IS-KO-AMR genes to the corresponding antibiotic, while blue bars indicate the opposite. **C)** Scheme of the pipeline followed to analyze the association between plasmid-borne IS elements and disruptions of KO-AMR genes. In this case, we employed the complete genomes available from the Bacterial and Viral Bioinformatics Resource Center (BV-BRC) as we could infer the genomic location of the IS. **D)** Pearson correlation between the number of plasmids encoded in the strains screened (x-axis) and the fraction of genomes with IS-KO-AMR genes (y-axis). We selected a threshold for the number of plasmids per strain which represented at least 95% of the database (n plasmids = 0-5, determined by quantile test). **E)** Results of analyzing the presence of plasmid-encoded ISs in strains carrying IS-KO-AMR gene mediated by the same IS. Y-axis indicates the Bonferroni adjusted p-value in log10 scale of the Chi square. X-axis indicates the IS element for which each statistical significance is represented. Filled points correspond to those IS which were found over enriched in plasmids in strains which showed KO-AMR inactivated by the same IS (indicating a positive relationship between carrying an IS encoded in a plasmid and a chromosomal gene inactivation mediated by the same IS).

### Plasmid-mediated gene inactivation is a generalised AMR mechanism

Our experimental observations are limited to pOXA-48 and the IS1. However, IS elements are overrepresented in plasmids^42^, and we therefore hypothesized that plasmids could generally promote AMR evolution through IS-mediated gene inactivation beyond our experimental model. To further investigate the potential prevalence and relevance of this phenomenon, we screened genomes from public repositories.

We first investigated the general contribution of IS elements to AMR evolution through gene inactivation. By combining the hits in MEGARes v.3.0 database^51^ and our own experimental targets, we identified a group of genes whose inactivation leads to AMR (KO-AMR genes, n = 54, **S. Table 4**). We screened all complete bacterial genomes available at the Bacterial and Viral Bioinformatics Resource Center (BV-BRC, n = 42,085) and detected that 1.82% of the bacterial strains presented IS-mediated knock-outs in KO-AMR genes (IS-KO-AMR, n = 766; **S. Fig. 4**). Most IS-KO-AMR genes were found in strains of the Enterobacterales (582/8129; ∼7.2%) and Pseudomonadales (178/1195; ∼14.9%) orders (**S. Fig. 4**). To confirm whether IS-KO-AMR genes were associated with a resistance phenotype, we screened the Enterobacterales genomes with available AMR-phenotype metadata at the Pathogen Detection Database of the NCBI (n = 19,177; **Fig. 3A**). As expected, we observed a significantly higher fraction of strains with a resistance level above the clinical threshold for its cognate antibiotic among those with an IS-KO-AMR gene (median fold-change = 3.74, p < 0.05, Fisher’s exact test after Bonferroni-Holm adjustment, **Fig. 3B, S**. **Table 4**). These results strongly suggest that IS-mediated gene inactivation plays a significant role in AMR evolution, particularly in Enterobactaerales. In fact, an overall analysis of genes and gene families inactivated by ISs elements (not only KO-AMR genes), revealed that most of the targets are likely associated with AMR phenotypes (**S. Fig. 4 C,D**).

**Figure 4.**
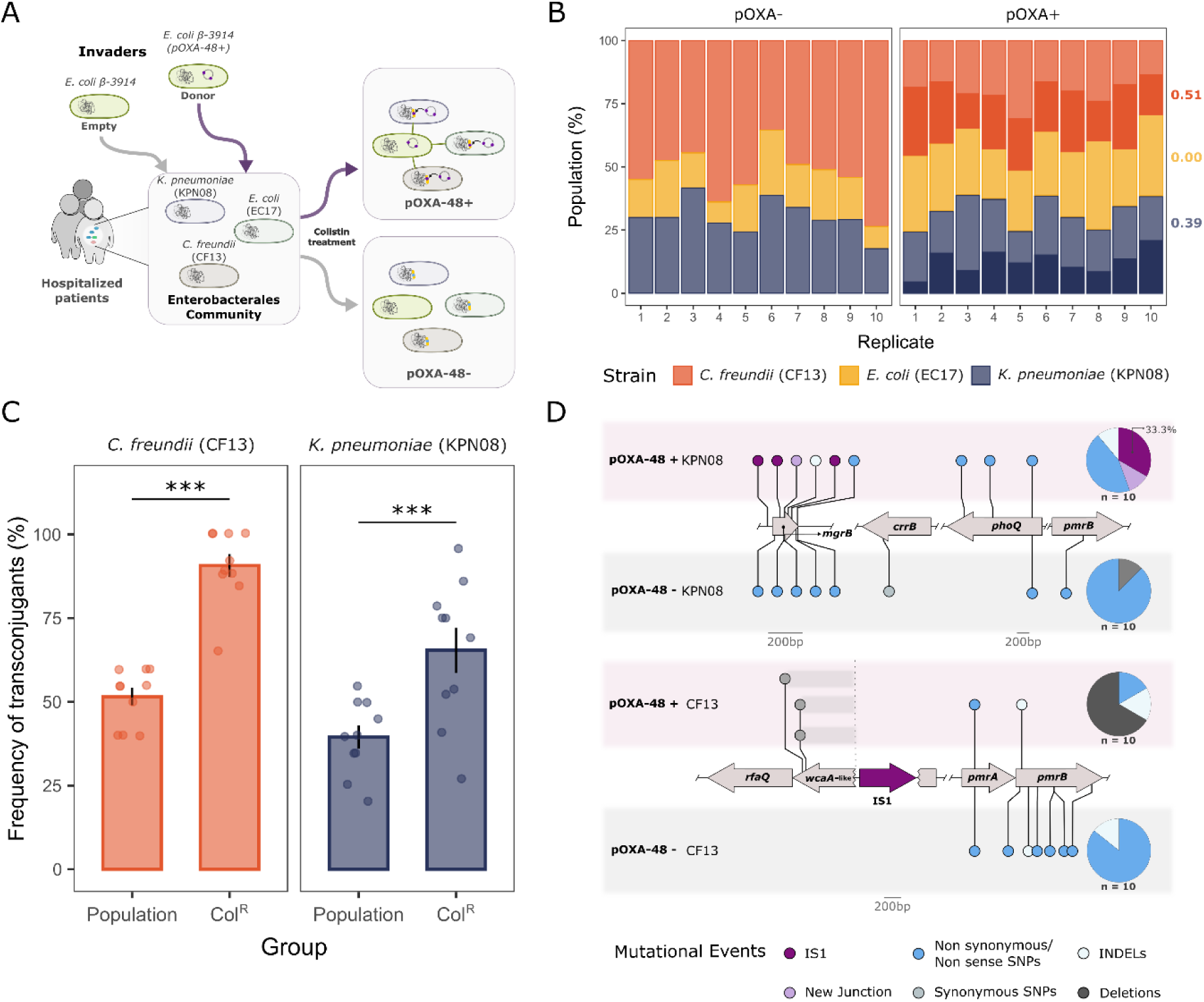
pOXA-48 community invasion promotes IS-mediated AMR. **A)** Schematic representation of pOXA-48 invasion of a community formed by three representative Enterobacterales commonly found in the gut microbiota of hospitalised patients (*K. pneumoniae* KPN08 [ST377], *E. coli* EC17 [ST131] and *C. freundii* CF13 [ST22]). We used the *E. coli* 𝛽-3914 strain carrying pOXA-48 to promote plasmid invasion in the communities (n=10), and pOXA-48-free 𝛽-3914 in the control communities (n=10). After the overnight invasion of the community, we plated communities in colistin and strain-selecting antibiotics. **B)** Community composition (%) per replicate population after counterselection of the empty *E. coli* 𝛽-3914 strain (left) and the pOXA-48-carrying *E. coli* 𝛽-3914 strain (right). The proportion of transconjugants for each strain is represented by darker bars. Values on the right of the plot indicate the mean frequency of transconjugants for each strain considering all the replicates (n = 10). Each color represents a given strain. **C)** Frequency of pOXA-48 transconjugants for either the whole population (left bar) or the colistin resistant group (right bar). We observed a significant increase in the frequency of transconjugants both in *C. freundii* and *K. pneumoniae* colistin resistant subpopulations when compared to the whole populations, as indicated by the asterisks (n = 10; p < 0.05; chi-squared test after Bonferroni adjustment). **D)** Mutations detected in the colistin resistant mutants isolated from the community. Mutations accumulated in the clones with no pOXA-48 are represented on the lower part of each strain panel (grey background), while mutations accumulated in the pOXA-48 transconjugants (purple background) are represented at the top of each strain panel. Pie charts represent the different types of mutations accumulated, as indicated in the legend. The number of strains sequenced for each condition is summarized in the right side of the panel, under each pie chart. Deletions putatively mediated by the IS1 are represented as grey bars covering the deleted genomic region. All genes of interest are plotted in order according to the reference genome, maintaining their relative positions while omitting the genetic regions between them.

Next, we analysed whether the IS-KO-AMR could be caused by plasmid-borne IS elements. To that end, we focused on the complete genomes from the BV-BRC (**Fig. 3C**), as we could determine the genomic location of the IS elements (see Methods). First, we detected a strong positive correlation between the fraction of genomes with IS-KO-AMR genes and the number of plasmids they carried (**Fig. 3D**; Pearson correlation R^2^ = 0.94, p < 0.05). Moreover, our results showed a significant association between carrying an IS-KO-AMR gene and the presence of a plasmid with that same IS element across the strains tested for 49 out of the 83 different IS elements (**Fig. 3E, S**. **Table 4**; Chi square test after Bonferroni adjustment, p < 0.05). These results support the notion that plasmids promote AMR evolution through IS-mediated gene inactivation.

Finally, we investigated what plasmid types could play a more relevant role in producing IS-KO-AMR genes. We observed that particular replicons, especially multi-replicon plasmids carrying either IncFII or IncFIB, were overrepresented in samples with IS-KO-AMR genes (**S. Fig. 5A**; p < 0.05 Fisher’s exact test after Bonferroni adjustment). Moreover, we found that these multi-replicon plasmids carrying IS elements from the IS5 and IS1 families were significantly enriched in strains with IS-KO-AMR genes disrupted by the same IS elements (**S. Fig. 5B**). These results strongly suggest that plasmid encoding IS elements play a major role in the emergence of AMR beyond our experimental model.

**Figure 5.**
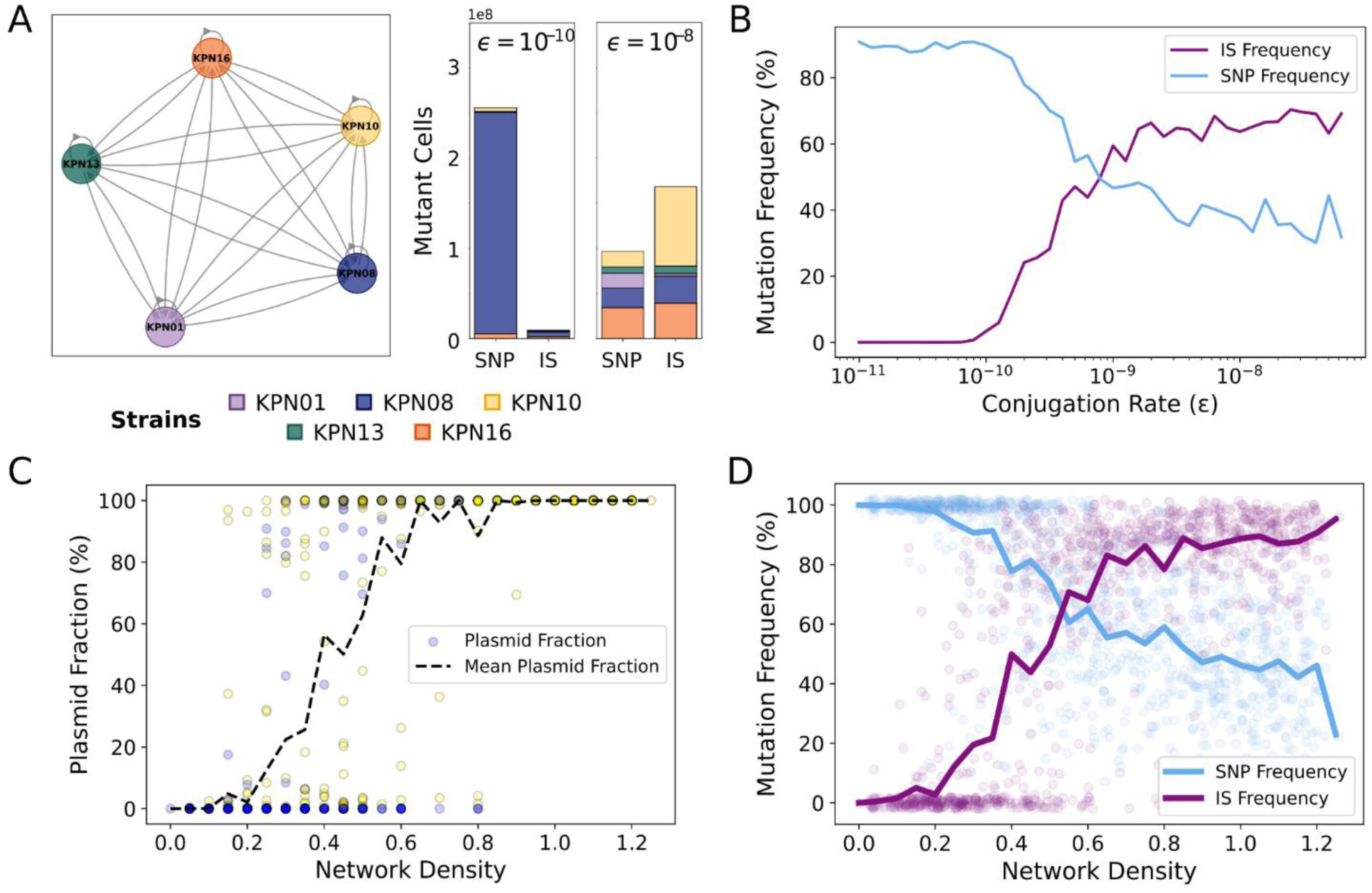
Simulations of plasmid transfer and mutation dynamics in multi-strain bacterial communities. **A)** Network representation of plasmid transfer between bacterial strains. Each node corresponds to a different strain, and directed edges indicate potential conjugation events. The conjugation rate (ε) modulates the likelihood of plasmid transfer along each edge. On the right, simulation snapshots showing the spatial distribution of mutants assuming an initial population of 10^8^ cells under two conjugation regimes, from left to right: ε = 10^-10^ and ε = 10^-8^. At lower conjugation rates (ε = 10^-10^), mutations arise primarily as SNPs. As ε increases, the mutational spectrum shifts, resulting in a higher frequency of IS-mediated mutations. Strain identity is indicated by color. **B)** Mutation frequency as a function of conjugation rate, averaged across simulations, as conjugation rate increases IS-mediated mutations also increase. **C)** Fraction of plasmid-bearing cells as a function of network density. The color of each point corresponds to the dominant strain (strain with the highest relative abundance). At higher network densities, plasmid-bearing cells dominate the population. **D)** Mutation frequency as a function of network density. Colored dots represent individual simulations; solid lines represent the average per network density. Blue corresponds to SNP mutations, and purple to IS-mediated mutations. Increased network connectivity correlates with a higher frequency of IS-mediated mutations.

### pOXA-48 invasion of a simple microbial community promotes AMR

Conjugative plasmids are the main vehicle for the dissemination of AMR genes in natural bacterial communities^52^. For example, we have previously shown that virtually every time a pOXA-48-carrying bacteria colonises the gut microbiota of a hospitalised patient, the plasmid conjugates between different strains^25^. Our new results make the tantalizing suggestion that by disseminating in bacterial communities, pOXA-48 may not only spread carbapenem resistance, but also promote the acquisition of further AMR mechanisms through IS1 transposition.

To test this hypothesis, we set up a simple community of three Enterobacterales strains isolated from the gut microbiota of hospitalised patients (from the R-GNOSIS collection^53^). These strains belonged to three different species, *Escherichia coli* (EC17), *K. pneumoniae* (KPN08) and *Citrobacter freundii* (CF13), and presented very different pOXA-48 reception frequencies by conjugation^54^. We selected *E. coli* 𝛽-3914 as the plasmid-carrying invader strain, because thanks to its counterselectable phenotype (auxotrophy for diaminopimelic acid)^55^, we could eventually eliminate it from the population and focus on the effects of pOXA-48 on the recipient strains. We performed parallel experiments adding the empty 𝛽-3914, or the pOXA-48-carrying 𝛽-3914 to the three-member communities (n = 10 biological replicates per treatment; **Fig. 4A**). After an overnight culture on solid agar, we analysed pOXA-48 conjugation frequencies and the frequency of COL resistant mutants in the different community members (see Methods). In line with our previous studies^54^, we observed marked differences in the conjugation frequencies among the community members, with *C. freundii* CF13 and *K. pneumoniae* KPN08 receiving the plasmid at high frequencies and *E. coli* EC17 producing no transconjugants (**Fig 4B**).

Contrary to our initial expectation, we did not observe a significant increase in COL resistance frequency in the communities invaded by pOXA-48 (t-test, n = 10, p > 0.05 after Bonferroni adjustment, **S. Fig. 6**). However, when we analysed the COL resistant subpopulations in the pOXA-48 treatment, we observed a significant association between pOXA-48 carriage and COL resistance in *C. freundii* and *K. pneumoniae* (**Fig. 4C**). To confirm the possibility that pOXA-48 acquisition facilitated COL resistance evolution in transconjugants, we sequenced the entire genome of a COL resistant clone from each treatment per independent replicate for *C. freundii* and *K. pneumoniae* (n = 40, 10 pOXA-48-free clones and 10 pOXA-48-carrying clones per strain). As expected, results revealed that pOXA-48 presence was associated with a higher frequency of IS1-mediated inactivations in *mgrB* leading to COL resistance in *K. pneumoniae* KPN08 (**Fig. 4D, S**. **Table 3**). In *C. freundii* CF13, we observed several SNPs or small deletions in COL resistance targets affecting the lipid A (*pmrAB, phoQ*) in both treatments^56^. Interestingly, we also found three independent cases of parallel evolution mediated by deletions in the LPS biosynthesis gene cluster specifically associated with pOXA-48 presence. These deletions were similar in the three clones, removing 600-700 nucleotides of the 3’ end of the *wcaA*-like glycosyltransferase gene. Crucially these deletions occurred immediately upstream of the only IS1 element encoded in the chromosome of this strain, which is inserted in the *wcaA*-like gene. Importantly, the genetic signatures of these events suggested that they may be mediated by the IS1, since the deletions started at one of the inverted repeat sequences (IR_L_) used to initiate transposition by the IS1 transposase InsAB. IS1 elements are known to produce deletions at the termini of the integrated element, while maintaining the element intact^57,58^. This process, known as IS1-mediated type II deletions, is independent of the cellular RecA system and does not involve DNA sequence homology. Our results strongly suggest that the pOXA-48-encoded IS1 elements, which are highly active^45^, could be responsible for these deletions. We provide a detailed analysis of these deletions in **S. Fig 7**.

### Computational model of plasmid invasion and IS-mediated AMR evolution

Our results with pOXA-48 suggested that conjugative plasmids can promote AMR evolution through gene inactivation while disseminating in bacterial communities. To explore this possibility beyond our experimental system, we built a computational model reproducing plasmid invasion of a bacterial community under different conjugation frequencies and antibiotic pressures. The model uses a stochastic Gillespie algorithm that explicitly simulates bacterial growth and antibiotic-induced death, plasmid dynamics including segregation and conjugation, and evolutionary events such as point mutations and IS transposition (Methods). This framework captures the complexity of multiclonal population dynamics and allows us to track evolutionary changes under antibiotic pressure. We estimated the model parameters directly from the *K. pneumoniae* clinical strains used in this study. Specifically, we estimated experimental growth, antibiotic susceptibility data (**S. Table. 1; S. Fig. 8**) and mutation rate measurements (from the mutation rates determinations, differentiating between SNPs and IS transposition, **S. Fig. 9**). The model tracks the accumulation of ISs and SNPs in plasmid-free and plasmid-bearing cells as they grow and compete for resources.

First, we simulated invasion experiments in multiclonal bacterial communities composed of five plasmid-free strains. At the start of each simulation, a small fraction of plasmid-bearing cells was introduced into the population. We implemented a serial transfer protocol to track evolutionary dynamics over time: populations were diluted daily, and resources and antibiotic concentrations were reset at each transfer. Plasmid transfer between strains was represented as a network, where edges define the potential for conjugation (**Fig. 5A**). In the complete network, all strains could exchange plasmids. As we gradually increased the conjugation rate in the complete network, we observed a sharp transition in plasmid prevalence: the population remained mostly plasmid-free at low transfer rates, but above a critical threshold, plasmids spread rapidly and reached fixation (**S. Fig. 10A**). Before this threshold, plasmid-free cells dominated the community; beyond it, multiple plasmid-bearing strains coexisted (**Fig. 5A; S. Fig. 10B**). This transition also shifted the AMR mutational spectrum: SNPs were more frequent when plasmids were rare, whereas IS transpositions became dominant after plasmid spread (**Fig. 5B**). Importantly, while this critical level of conjugation was high (ε **∼** 10^-9^), many clinically relevant AMR plasmids conjugate at this rate^59^.

To further explore how the structure of the transfer network influences evolutionary outcomes, we generated 1000 random plasmid transfer matrices by randomly removing edges, sampling a wide range of network densities while keeping the conjugation rate constant (ε = 10^-9^, which is similar to pOXA-48 conjugation rate, **S. Fig. 11**). Both plasmid fraction and average number of mutations per cell increased with network density (**Fig. 5C–D**). At low densities, SNPs were more common. In contrast, at higher densities, ISs became the predominant mutation type (**Fig. 5D**). The results from this computational model confirmed that conjugative plasmids can promote AMR evolution through IS-mediated gene inactivation under realistic parameters of plasmid transfer rate and community structure.

## Discussion

In this study, we first demonstrated that plasmid pOXA-48 promotes AMR evolution through the IS1-mediated inactivation of chromosomal genes (**Figs. 1, 2 and 4**). Next, we expanded this observation beyond our experimental system through genome-wide bioinformatic screens and computational modelling (**Figs. 3 and 5**). Analyzing genome databases, we were able to link the presence of plasmid-encoded IS elements with the disruption of chromosomal genes leading to AMR for multiple antibiotics and across diverse bacterial taxa (**Fig. 3**; **S. Fig. 3**). Interestingly, some of the most common Enterobacterales plasmid types (F-like) and IS families (IS5 [ISKpn26; IS903B] and IS1 [ISKpn14; IS1A]) were enriched in genomes with IS-mediated inactivations leading to AMR. This observation goes in line with results from previous experimental observations of F-like plasmids and IS5-like and IS1-like promoting AMR evolution through this route^60–62^. These results strongly suggest that plasmid-mediated IS gene inactivation plays a central role in AMR evolution in Enterobacterales. Finally, we developed a computational model simulating the invasion of a bacterial community by a conjugative plasmid harbouring IS elements. This model allowed us to explore a range of realistic parameters of plasmid dissemination, demonstrating that plasmid invasion can generally promote the IS-driven evolution of AMR (**Fig. 5**). Taken together, these results indicate that conjugative plasmids not only fuel AMR evolution through the dissemination of resistance genes, but also through IS-mediated inactivation of chromosomal genes.

Our study has important clinical implications. Carbapenem resistant Enterobacterales (CRE) are among the most threatening antibiotic-resistant bacterial pathogens due to their ability to surpass nearly every available therapy. The use of COL as a last-resort antibiotic has been a useful strategy to treat carbapenem-resistant infections in clinical settings. However, recent studies have evidenced a concerning rise of COL resistance in CRE^37,40,41^, critically limiting the treatment options. Importantly, the disruption of chromosomal genes by IS stands out as one of the main routes of acquisition of COL resistance^39,63^. The results of this study can explain the accelerated rise of combined resistance phenotypes to these two last-resort antibiotics^60,61^. Considering the ubiquitous distribution of bacterial plasmids, and the abundance of studies reporting AMR to several antibiotics through IS-mediated inactivation -including carbapenems, cephalosporins and aminoglycosides^31^-plasmids might play a critical role in AMR evolution through this new route.

IS are major evolutionary drivers in bacteria^28,64^, and plasmids -especially conjugative plasmids^65^-show an over enrichment in IS elements^42^. This fact supports the idea of a synergistic relationship between both MGE. Namely, conjugative plasmids can increase the frequency (i.e. fitness) of IS elements by allowing them to invade new bacterial hosts, while IS elements could indirectly promote plasmid fitness through second-order selection^66^, via adaptive mutations in the bacterial chromosome. As a result, plasmids can act as dual platforms for gene dissemination and gene inactivation, fostering rapid bacterial adaptation^67^. Indeed, it has been shown that machine learning models which include resistance genes together with IS presence in the genomes of *Acinetobacter baumannii* significantly improve AMR phenotypic prediction^32^. Our model can serve as the basis for future work exploring plasmid-encoded IS driven evolution using plasmids/ISs with diverse conjugation/transposition rates. Finally, a key potential advantage associated with IS-mediated inactivations is that they not only occur at a higher rate than SNP-mediated inactivations, but they also revert at a higher rate^68,69^. The high inactivation/reversion rates mediated by the insertion/excision of IS elements will be particularly useful in fluctuating environments, such as those faced by bacteria during antibiotic treatments.

Another important result of our study is the ability of pOXA-48 to promote IS-mediated AMR while invading a bacterial community composed of clinical Enterobacterales recovered from hospitalised patients. The experiments showed how, even if one of the community members does not accept the plasmid, acting as a roadblock for conjugation, pOXA-48 can still successfully invade the remaining members (*K. pneumoniae* KPN08 and *C. freundii* CF13). Importantly, pOXA-48 promoted COL resistance acquisition in this process. In *K. pneumoniae* KPN08, pOXA-48 induced once again IS1-mediated inactivation of chromosomal genes leading to COL resistance. In *C. freundii* we observed a puzzling result. While for some clones COL resistance could be explained by mutations in classic targets (*pmrAB*), one particular deletion in the LPS gene cluster was specifically linked to COL resistance when pOXA-48 was present. Interestingly, our analyses suggested that the pOXA-48-encoded IS1 elements may be responsible for these deletions, linking IS-mediated AMR acquisition with yet another mechanism: IS1-mediated type II deletions. These results provide an extra layer of complexity in our understanding of the interplay between MGEs and AMR evolution.

One important limitation of our work is that we mainly focused on the order Enterobacterales. Due to their clinical relevance, public databases show an overrepresentation of Enterobacterales genomes^70^ (**S. Fig 3**), which results in phylogenetically biased datasets. Nonetheless, both our results (**Figs. 3 and 5**) and recent evidence^32,33^ indicate that this mechanism also occurs in other bacteria such as *A. baumannii* or *E. faecium*. Given the considerable abundance and widespread nature of both plasmids and IS in prokaryotic genomes^71,72^, we anticipate that plasmids act as main drivers of IS-mediated evolution of AMR in many bacterial pathogens.

## Methods

### Bacterial strains

All the clinical strains included in this work belong to the R-GNOSIS collection, obtained as part of a surveillance screening program for detecting extended-spectrum ß-lactamases or carbapenemase-producing Enterobacterales in hospitalized patients at the Hospital Universitario Ramón y Cajal^53^ (Madrid, 2014-206), approved by the Hospital Ethics Committee (ref. no, 251/13). The strains selected in the initial screen include 20 *K. pneumoniae* (ST15, ST377, ST432 and ST1427), one *E. coli* (ST131) and one *C. freundii* (ST22). For the community assay, we also used a counterselectable diaminopimelic acid auxotrophic laboratory strain, the *E. coli* 𝛽-3914^55^. A complete description of the strains can be found in **S. Table 1**.

### Construction of pOXA-48 ΔΔIS1

To confirm whether the results were caused by the IS1s encoded in pOXA-48 or by the presence of the plasmid *per se*, we constructed a version of pOXA-48 lacking both IS1s using lambda-red recombination. We amplified the recombination cassette with extremes matching the beginning and end of IS1 (9829-10104), and a KAN gene flanked by FRT sides from pKD4, as described in Datsenko *et al.*^73^ Similarly, we amplified the recombination cassette with extremes matching the beginning and end of IS1 (32412-32789), and a CMP gene flanked by FRT sites similarly from pKD3. The primers used for construction of the cassette and validation of the double mutant construct can be found in **S. Table 5**. ***Fluctuation assays***

To study the phenotypic resistance mutation rate against antibiotics, we performed fluctuation assays^46^. We conducted fluctuation assays using the following strains with and without pOXA-48: KPN01, KPN08, KPN10, KPN13 and KPN16. In addition, we conjugated the pOXA-48ΔΔIS1 into the KPN08 strain as a control, and included this strain in the assays. We inoculated each strain from freezer stock onto LB agar plates and incubated them at 37°C for 18 hours. We selected 35 independent colonies for each strain and cultured them overnight in 2 mL of LB broth with continuous shaking. Then, we inoculated 15 μL of each overnight culture either in a LB agar plate with 2 mg/mL of COL (Sigma-Aldrich, EEUU) or 20 mg/mL RIF (Sanofi, Spain). To account for the number of viable cells we also inoculated 15 μL of each overnight culture in LB agar plates with no antibiotic. We counted the number of resistant mutants in the dilution 10^-2^ for antibiotic-containing plates and the number of viable cells in the dilution 10^-7^ for the LB plates. Finally, we selected 10 colonies from independent cultures for each strain in the presence of COL. This was achieved in all the samples and conditions except for our construction pOXA-48ΔΔIS1 (n = 9 colonies) and pOXA-48-carrying KPN13 (n = 8 colonies). We sequenced the whole genome of 105 samples (see *Whole Genome Sequencing* section).

### High-throughput antibiotic susceptibility testing

To study the role of pOXA-48 in the acquisition of resistance to other antibiotics, we followed an experimental approach which allows for high-throughput analysis of resistance to multiple antibiotic concentrations. We performed the assay for two of the strains (KPN08 and KPN16) with and without pOXA-48. For KPN08 we tested COL, CIP, CMP and RIF and for KPN16 COL, CIP, KAN and RIF (due to their different susceptibility patterns). Before the experiment, we determined the MICs of the strains for their respective antibiotics (see *Extended Methods*). We selected 47 colonies from an overnight culture plated on LB-agar without antibiotic for each isogenic version of the strains. We inoculated each colony in 200 μL of LB in 96 well plates (distributed in checkerboard pattern to avoid contamination). We incubated the plate for 22 hours at 37°C with constant shaking (Synergy HTX Multi-Mode Reader; BioTek Instruments; 100 rpm). We transferred 2 μL of each culture into a new plate with 198μL of LB with increasing antibiotics concentration per well. After overnight culture in the same conditions, we measured the OD600 for each of the cultures for both versions of the strain to analyze the number of surviving replicates in the increasing concentrations of antibiotics. We calculated the median OD600 for the positive controls obtained from the same overnight culture. We classified the samples as surviving if their growth was over 10% of the median positive control (i.e. IC90) or not surviving if it was below. Finally, we sequenced the whole genome of 111 samples, including all strains with and without pOXA-48 tested against the different antibiotics. The samples were selected from the highest antibiotic concentration that showed growth for both conditions. To reduce the number of sequencing reactions, we mixed two clones and sequenced them as a single population (see *Whole Genome Sequencing* section).

### Community experiment

#### Study design and strain selection

To recreate the invasion of a community with plasmid pOXA-48, we used three strains isolated from the gut microbiota of hospitalised patients: *Citrobacter freundii* ST22 (CF13^45^), *Klebsiella pneumoniae* ST11 (KPN06), and *Escherichia coli* ST131 (EC17). The fourth member of the community was either a pOXA-48-carrying or a pOXA-48-free *Escherichia coli* ß3914 which is auxotrophic for diaminopimelic acid (DAP). First, we determined the antibiotic susceptibility of each community member to colistin. We inoculated 2mL of LB from the frozen stock of each member and incubated them overnight (37°C, shaking 250 rpm). The next day, we serially diluted the cultures in 0.9% NaCl and plated 10 µL from 1:10 to 1:10^7^ dilutions onto LB agar plates supplemented with 1 or 2 µg/mL of colistin. Similarly, we determined the antibiotic susceptibility of each community member to Zeocin (ZEO), Apramycin (APR) and Hygromycin B (HYG) following the same procedure as described before and plating 100 µL of each culture onto LB agar plates with each antibiotic at different concentrations. After an overnight incubation at 37°C, we determined the susceptibility levels of each member.

#### Marker plasmid construction

We used three different non-transmissible plasmids derived from pBGC in order to individually select each member of the community. We used pBGA (APR^R^), constructed in a previous work^50^, pBGZ (ZEO^R^) and pBGHB (HYG^R^). For pBGZ and pBGHB, we respectively replaced the *catA1* gene from pBGC with the *SH ble* gene from pKM496 (Addgene #109301) or the *aph(7’’)-Ia* gene from pKM464 (Addgene #108322) using Gibson assembly (NEBuilder HiFi DNA Assembly, USA). We validated plasmid sequences by long-read sequencing (Plasmidsaurus, USA) after plasmid extraction (NucleoSpin Plasmid, Macherey-Nagel, USA).

#### Introduction of marker plasmid

We prepared competent cells for each community member by washing 5 mL of overnight LB cultures (grown at 37°C and 250 rpm) with 50 mL of sterile pure water (Thermo Fisher Scientific). We introduced each plasmid by electroporation (MicroPulser Electroporator; Bio-Rad Laboratories). To select for each plasmid in each strain, we used the following antibiotics: ZEO 200 mg/L for PF_KPN08+pBGZ, HYG 120 mg/L for CF13_pOXA-48-free+pBGHB, and APR 50 mg/L for PF_EC17+pBGA. Plasmid presence was confirmed by PCR (see **S. Table 6**). One colony per community member was selected to start LB cultures supplemented with each antibiotic and grown overnight (37°C and 250 rpm) to be stored.

#### Community experiment

Prior to this experiment, we evaluated and discarded plasmid loss and possible antagonistic interactions (see *Extended methods*). For the community experiment, we inoculated 2 mL LB cultures of each member from glycerol stocks and cultured them overnight (n = 10 for each pOXA-48/pOXA-48-free community). We mixed 300 µL of each culture into a single tube and plated 50 µL of the mixture onto an LB-agar plate supplemented with 0.3 mM DAP. We incubated the plate for 24 hours at 37°C. Next, we recovered the biomass and resuspended it in 5 mL of LB supplemented with 0.3 mM DAP. We diluted the mix (1:100) in 5 mL of fresh LB with 0.3 mM DAP and incubated it overnight (37°C, 250 rpm). The next day, we serially diluted the cultures (from 1:10 to 1:10^10^) in sterile 0.9% NaCl and we plated the dilutions on LB-agar plates supplemented with each antibiotic. We used dilutions 1:100 to 1:10^9^ to seed LB agar plates supplemented with each selective antibiotic and 0.3 mM DAP, as well as antibiotic-free LB-agar plates. We used dilutions 1:1 to 1:100 to seed 1 µg/mL colistin plates, to determine the COL mutation frequency in each community. Additionally, COL plus strain-selective antibiotic plates were also seeded, to analyse the COL mutation frequency for each strain. After incubating the plates overnight at 37°C, we determined the colony number in each plate. Finally, we selected a COL resistant clone from each species per independent replicate to perform whole genome sequencing (N = 40, 20 pOXA-48-free clones and 20 pOXA-48-carrying clones). To reduce the number of sequencing reactions, we mixed two clones from different replicates and sequenced them as a single population (see *Whole Genome Sequencing* section).

### Whole genome sequencing

#### Illumina short-read sequencing

We extracted the genomic DNA using the Wizard Genomic DNA Purification Kit (Promega Corporation). To analyze the mutations that generated resistant phenotype, we isolated and sequenced 256 resistant clones from the fluctuation (n = 105), high-throughput susceptibility (n = 111) and community (n = 40) assays. The short-read sequencing of these samples was performed at SeqCoast Genomics (New Hampshire, USA) on an Illumina NextSeq200 platform, producing 2x150bp paired reads (200 Mbp per sample; >30x coverage per clone).

#### Oxford Nanopore Technologies long-read sequencing

To get closed assembled reference genomes of the clinical strains (**S. Table 1**), we sequenced them using Oxford Nanopore Technologies (ONT). We extracted the genomic DNA as explained in the above section. We prepared the genomic libraries following the Nanopore protocol for native barcoding genomic DNA (EXP-NBD114.24 kit). We quantified the DNA using a Qubit Flex Fluorometer (Thermo Fisher Scientific). To perform the DNA repair and end-prep steps, we used the NEBNext Ligation Sequencing Kit (Oxford Nanopore Technologies), AMPure XP beads (Beckman Coulter/Agencourt AMPure XP) and a HulaMixer (Thermo Fisher Scientific). For the native barcode ligation step, we used the EXP-NBD114.24 kit (Oxford Nanopore Technologies) and the Blunt/TA Ligase Master Mix (New England Biolabs). Then, we performed the adaptor ligation and clean-up steps with the NEBNext Quick Ligation Reaction Buffer (New England Biolabs), NEBNext Ligation Sequencing Kit (New England Biolabs). We carried out the priming and loading of the flow cell steps, in which we used the Flow Cell Priming Kit (EXP-FLP004) (Oxford Nanopore Technologies) and sequenced the samples using R10.4.1 Flow Cells (FLO-MIN114) in a MinION Mk1B.

### Characterization of genomic mutations

#### Assembly of reference genomes of clinical strains

We assembled the genomes of the stocked isolated strains carrying pOXA-48, to later use them as reference genomes. We performed the hybrid assembly of both long (ONT) and short (Illumina) reads using Unicycler v.0.5.0. We confirmed the assembly completeness using Bandage v.0.9. We then annotated the assembled genomes with Bakta v.1.9.3.

#### Quality control of Illumina reads

We carried out a quality control of the raw Illumina reads using FastQC v.0.12.1 and merged all the reports with MultiQC v.1.27.1. We trimmed the reads using Trim Galore v.0.6.10, adapting the parameters to trim low-quality ends, discard reads shorter than 50 bp and to trim nextera adaptors (-q 20 –length 50 –nextera). We performed a second quality control of the trimmed reads and used them for the following downstream analyses.

#### Variant calling

To detect mutational events in resistant mutants, we performed the variant calling of the Illumina short reads using breseq v.39.0.^74^ against the assembled reference genome of each strain. For the population WGS data obtained from the high-throughput and community assays, we used the ‘-p’ flag to detect polymorphic mutations. To analyze the results, we developed a Python script (Python v.3.8) to parse the HTML results given by breseq to obtain xlsx files (see ‘Code availability’) and simultaneously compare mutations present in the samples by merging their reports into an individual table per strain. For further information on the filtering criteria see *Extended methods*.

### Database analyses

#### Local reference construction

We built a database based on the ISfinder available data (https://github.com/thanhleviet/Isfinder-sequences; 2020-Oct, n = 5970). We downloaded the MEGARes database latest release (3.0.0; 2022-09-15; n = 8733) and built again a local reference database to look for disrupted AMR related genes (genes that confer AMR when disrupted), using ABRicate. Additionally, we included our experimental targets that confer AMR when disrupted, some of which were not present in the MEGARes database (*rbsB, araF, crrB, mgrB, oqxR, pmrB, qseC, sapB, sbmA, lepA, yebO* and *yobH;* n = 12).

#### Analysis of ISs impact on AMR phenotype

To study the impact of IS-mediated KOs on KO-AMR genes, we analyzed the genomes available at the NCBI Pathogen Detection Database (https://www.ncbi.nlm.nih.gov/pathogens/ast/<u>#</u>), which contains metadata of the AMR phenotype of the strains. We first retrieved the genome assemblies (last access January 2025) with metadata associated to the antibiotics we experimentally tested (CMP, CIP, KAN, and polymyxins—including COL and polymyxin E) plus fosfomycin and all the available beta-lactams. We discarded antibiotics with fewer than 100 total samples or fewer than 5 samples in either the resistant or susceptible phenotype. Based on these criteria, 21 antibiotics were filtered (final number of antibiotics analysed = 29; n = 19,162). To detect disrupted genes, we ran ABRicate on the genomes using both IS and AMR databases setting the minimum alignment coverage to 5% and the minimum nucleotide identity to 50% to facilitate the detection of both elements even if they were located at the end of a contig. We defined chromosomal AMR-related genes as disrupted when an IS element was detected adjacent to a gene mapped with partial coverage (<100%). Finally, we statistically assessed the differences in genomes showing KO-AMR genes between resistant and susceptible groups for each antibiotic (See *ISs-mediated KOs causing resistance phenotype* methods section).

#### Distribution of KO-AMR genes by plasmid-borne ISs

To study the generalized distribution of AMR targets of plasmid-encoded IS elements, we analyzed the complete genomes available at the BV-BRC (Bacterial and Viral Bioinformatics Resource Center; https://www.bv-brc.org/). We retrieved the samples filtering by genome completeness (complete) and quality (good), and analyzed them after removing duplicates and genomes which contained fragmented assemblies (n = 42,085; last accessed July 2024). Using the IS local database, we ran ABRicate with default parameters (minimum alignment coverage = 80%; minimum nucleotide identity = 80%). To detect disrupted target genes (i.e. partial alignments detecting both extremes of the KO AMR determinant), we set the minimum alignment coverage to 30% and the minimum nucleotide identity to 80% (as default, as nucleotide sequences are less conserved than protein sequences; however, the search is much faster using nucleotide query and subject sequences). Finally, we considered the IS-mediated KO-AMR genes, those for which we detected an IS element or a transposon inserted between both ends of the gene identified. Then, we statistically analyzed whether most of these IS elements were plasmid-borne (see *Plasmid-borne ISs causing KOs detected in databases* section in statistics methods part).

### Statistics

#### Phenotypic resistant mutation rate analysis

We inferred the phenotypic resistant mutation rate using the rSalvador package^47^ and compared it for each strain with and without pOXA-48 applying maximum likelihood ratio tests adjusted for multiple testing by the Bonferroni-Holm method. We applied the same reasoning for all the fluctuation assays included in this work, including the pOXA-48ΔΔIS1 test, in which case we compared the 3 genotypes of the experiment (pOXA-48-free, pOXA-48-carrying and pOXA-48ΔΔIS1-carrying).

#### Logistic regression models of strains survival

To statistically compare the survival of the strains on increasing concentrations of antibiotics during the high-throughput susceptibility assays between pOXA-48-free and pOXA-48-carrying pairs, we built a logistic regression model per antibiotic and strain. We first classified for each antibiotic concentration, strain genotype and antibiotic used, the samples in surviving or dead as explained in the High-throughput antibiotic susceptibility testing section. We then modeled the survival probability of the strain with or without pOXA-48 against each antibiotic used using a binomial distribution of the data. We fitted a binary logistic regression with a logit link function between the number of surviving samples and the antibiotic concentration for each strain and antibiotic. We then applied the predicted fitted regression to the observed data for each genotype and studied the statistical significance of both concentration and genotype on the survival of the strains. The formula can be defined as:

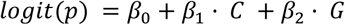

Where *p* is the survival probability of a given sample; *ß*_0_ is the intercept, *ß*_1_ is the coefficient for the antibiotic concentration (*C*) variable; and *ß*_2_ is the coefficient for the strain genotype (*G*) variable. Specifically, *p* is encoded as a binary outcome (1 = live, 0 = dead), *C* varies for each antibiotic and strain (see High-throughput antibiotic susceptibility testing methods section) and *G* is either pOXA-48- or pOXA-48+.

#### Plasmid-borne ISs causing KOs detected in databases

To infer whether the KO of chromosomal genes detected in the BV-BRC database were potentially associated with the IS encoded in plasmids present in each genome, we statistically analyzed the results obtained from the analysis indicated in the *Distribution of AMR determinants KOs by plasmid-borne ISs* section. Specifically, for each IS element, we performed a Fisher’s exact test. We constructed a 2×2 contingency table and classified the genomes based on the presence of plasmid-encoded IS elements and the occurrence of AMR gene disruption, getting 4 possibilities: **A** = number of genomes with plasmids encoding the IS element causing the chromosomal KO detected; **B** = number of genomes with plasmids encoding the IS element but not showing AMR-related KOs; **C** = number of genomes without plasmids encoding the IS-element but showing chromosomal AMR-related gene disrupted; **D** = number of genomes not showing neither AMR-related gene disrupted nor IS element encoded in a plasmid. After performing the exact Fisher’s tests, we adjusted the p-values using the Bonferroni-Hochberg method.

#### ISs-mediated KOs causing resistance phenotype

To analyze whether KO-AMR mediated by IS were enriched in resistant strains to multiple antibiotics (n = 29, see *Analysis of ISs impact on AMR phenotype*), we compared frequency of resistant strains by IS-mediated KO-AMR with the frequency of resistant strains to a certain antibiotic using Fisher’s exact tests after Bonferroni-Holm adjustment for multiple comparisons.

### Computational model

#### Stochastic Model of Plasmid Dynamics and Mutation Accumulation

We developed a stochastic, multi-species model to simulate the evolutionary dynamics of bacterial populations under selection. We modeled all events as discrete stochastic transitions, with propensities determined by strain-specific parameters. Growth was modeled using monod kinetics under a homogeneous environment, with resources equally accessible to all cells. We incorporated antibiotic pressure by modulating death rates via a saturating dose–response function. Plasmid segregation occurred stochastically at cell division, resulting in the plasmid’s loss and the cell’s reclassification into the plasmid-free subpopulation. We modeled conjugation as a mass-action process, with successful transfer dependent on the conjugation rate and the permissiveness of the recipient cell. The simulated community was composed of 5 species, each partitioned into plasmid-free and plasmid-bearing subpopulations. Mutation and transposition events occurred independently and irreversibly. We assumed all mutations affect phenotype, either conferring resistance or imposing a fitness cost. In the base case, we presumed a fully connected network in which all strains can transfer plasmids to all others at a uniform rate. We estimated model parameters independently for each bacterial strain using experimental data obtained from monoculture growth assays and mutation rate measurements. We used the quantification of events obtained from the whole-genome sequencing of replicates at the end of the fluctuation assays to calculate the SNP and IS mutation rates. For full technical specifications of the model see Extended Methods.

#### Simulating Evolutionary Dynamics in Serial Transfer Experiments

We ran simulations in discrete 24-hour cycles, representing daily growth phases. At the end of each cycle, populations were diluted by a fixed factor (1:100) to simulate serial transfer conditions, maintaining relative strain and plasmid frequencies while reducing absolute cell numbers. Resource and antibiotic concentrations were replenished to their initial values at each transfer, allowing for consistent environmental conditions across days. This setup allowed us to track plasmid dynamics and the accumulation of mutations over 45 simulated days. We performed two classes of *in silico* experiments. In the first, we fixed the network topology as a complete graph in which all strains could exchange plasmids with one another and we varied the conjugation rate (ε). In the second set of experiments, we fixed ε=10⁻^9^. We randomly removed edges from the complete graph to systematically vary network topology while preserving global connectivity. To quantify evolutionary outcomes, we tracked the population dynamics of each strain over time, including total cell counts, plasmid status, and the accumulation of SNPs and ISs in both plasmid-free and plasmid-bearing subpopulations.

## Code availability

The code developed for this project can be found in https://github.com/jorgEVOplasmids/MDR_ISs.

## Data availability

All the sequences generated for this project can be found at the Sequence Read Archive (SRA) repository of the National Center for Biotechnology Information (NCBI) under the BioProject ID PRJNA1294581.

## Contributions

J.S.-D., P.R.-F., J.D., A.S.-L. and A.S.M. conceptualized the study. J.S.-D., P.R.-F., J.D, A.S.-L. and A.S.M. designed the methodology. J.S.-D and P.R.-F. analysed the genomic data. J.D., S.M.-G., S.Q., M.V.-R., and C.C. performed the experiments. A.C.-V. contributed with tools, A.F.-H performed the mathematical model. J.S.-D., P.R.-F., J.D., A.S.-L. and A.S.M. analysed the data, prepared the original draft of the manuscript and reviewed and edited the manuscript. A.S.-L. and A.S.M. acquired the funding and supervised the study.

## Acknowledgements

We thank J.A. Escudero, J. Rodríguez Beltrán and J. Penadés for constructive comments on the manuscript. Work in the A.S.M. lab was supported by the European Research Council (ERC) under the European Union’s Horizon 2020 research and innovation programme (ERC StG grant no. 757440-PLASREVOLUTION), and under the European Union’s Horizon Europe research and innovation programme (ERC-2022-CoG Project 101086992-PLAS-FIGHTER), and by Project PCI2021-122062-2A funded by MICIU/AEI/10.13039/501100011033 and by the European Union NextGenerationEU/PRTR. The A.S.-L. lab acknowledges support from Project PID2023-152460NA-I00, funded by MICIU/AEI/10.13039/501100011033 and by ERDF/EU, the ‘La Caixa’ Foundation (ID 100010434) under project LCF/BQ/PR22/11920001; and by Grant RYC2022-037765-I, funded by MICIU/AEI/10.13039/501100011033 and by European Union NextGenerationEU/PRTR. A.CV. was funded by an EMBO postdoctoral fellowship (ALTF 322-2022). A.F.H. was funded by the sabbatical program DGAPA-PASPA, UNAM.

## Supplementary Figures

**Supplementary Figure 1.**
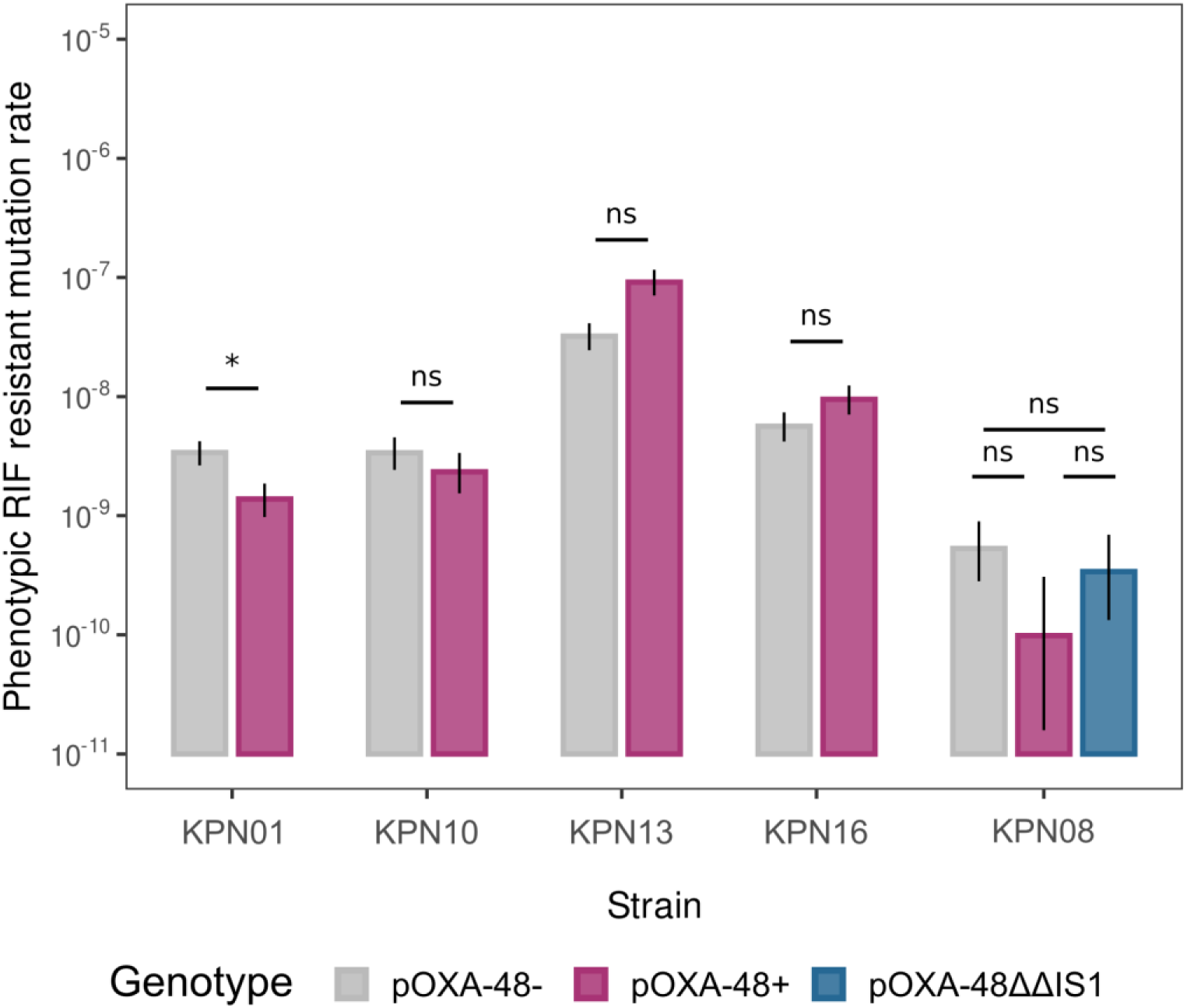
Mutation rate to rifampicin of isogenic clinical pairs carrying or not carrying pOXA-48. Phenotypic resistance mutation rate to rifampicin of the isogenic pairs of *K. pneumoniae* clinical strains either not carrying (grey) or carrying pOXA-48 (purple), and carrying the IS1-depleted version of pOXA-48 (pOXA-48ΔΔIS1). The black bars indicate the 95% confidence interval. Asterisks denote statistical significance in the mutation rate between pOXA-48- and pOXA-48+ for each pair based on maximum likelihood ratio tests (n = 35) after Bonferroni-Holm adjustment for multiple testing (p-value: 0 < *** < 0.001 < ** < 0.01 < * < 0.05 < ns).

**Supplementary Figure 2.**
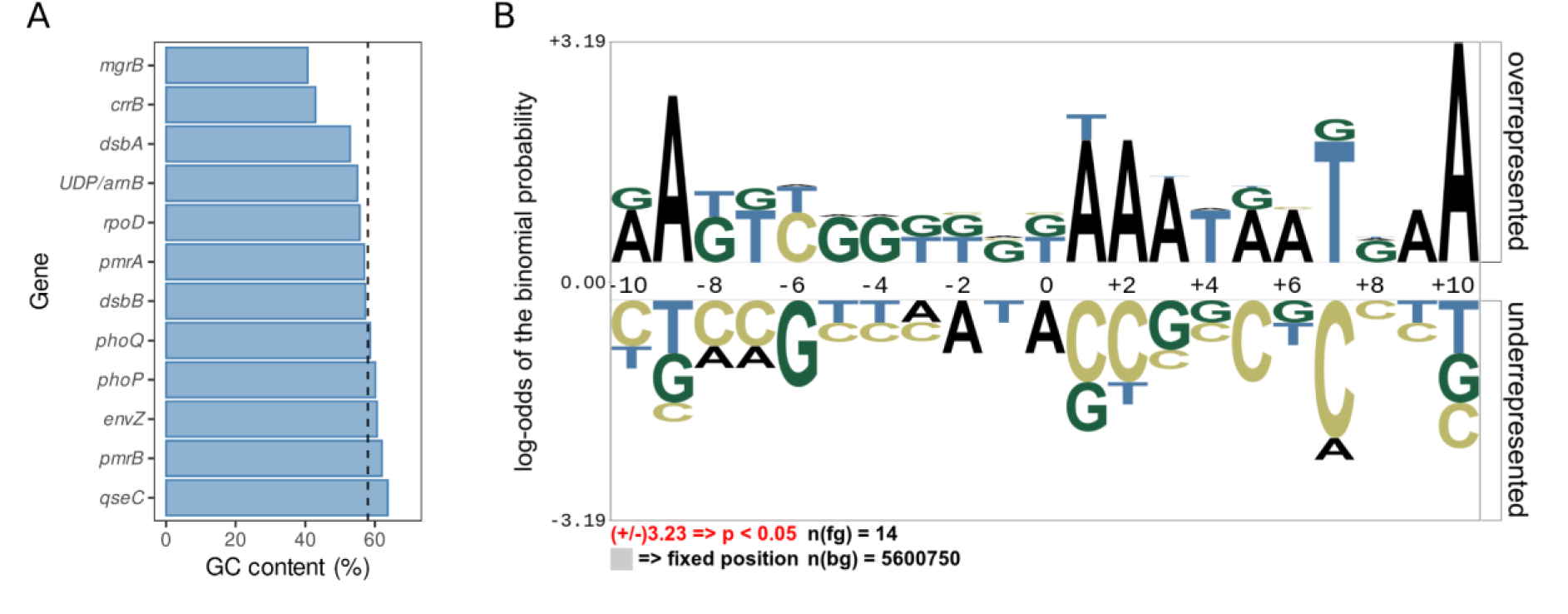
GC content of colistin resistant mutants target genes. **A)** Barplot of the GC content of the genes targeted during the fluctuation tests in colistin resistant mutants. Dashed black line indicates the mean GC content of the *K. pneumoniae* chromosome (∼57%). *mgrB*, the main target of IS1 insertions during the experiment, showed the lowest GC content. **B)** Sequence logo of the IS1 insertion sites. We observed an overrepresentation of the AAATAATGAA sequence in the +10 nucleotide positions following the insertion sites.

**Supplementary Figure 3.**
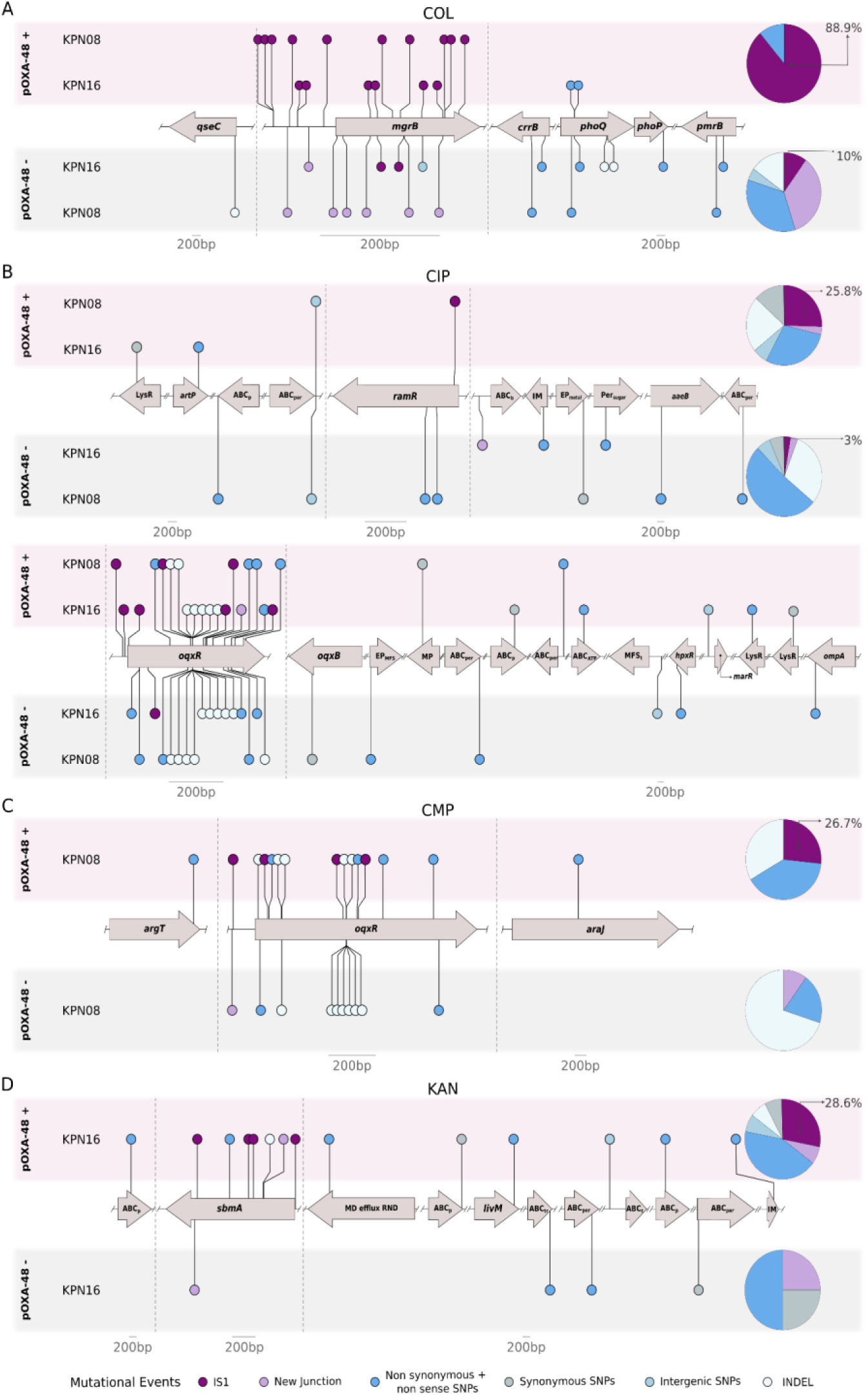
Genetic targets of mutants resistant to COL, CIP, CMP and KAN. All mutations detected in the common highest concentration for both conditions (pOXA-48 carrying and pOXA-48 free strains) in the evolutionary rescue assays for Colistin (**A**), Ciprofloxacin (**B**), Chloramphenicol (**C**) and Kanamycin (**D**). Mutations accumulated in absence of pOXA-48 are represented on the lower part of the panel (grey background) while those accumulated in presence of pOXA-48 are represented at the top of the panel (purple background). Pie charts of all the events are displayed on the right for each condition. New junctions mediated by IS1 elements are represented in dark purple while new junctions mediated by other elements are in light purple. Dark blue indicates protein-altering SNPs, light blue denotes intergenic SNPs, light grey represents synonymous SNPs, and white denotes INDELs. All genes are plotted in order according to the reference genome, maintaining their relative position. The most relevant genes with the highest number of events have been zoomed in, represented by the gray dashed line.

**Supplementary Figure 4.**
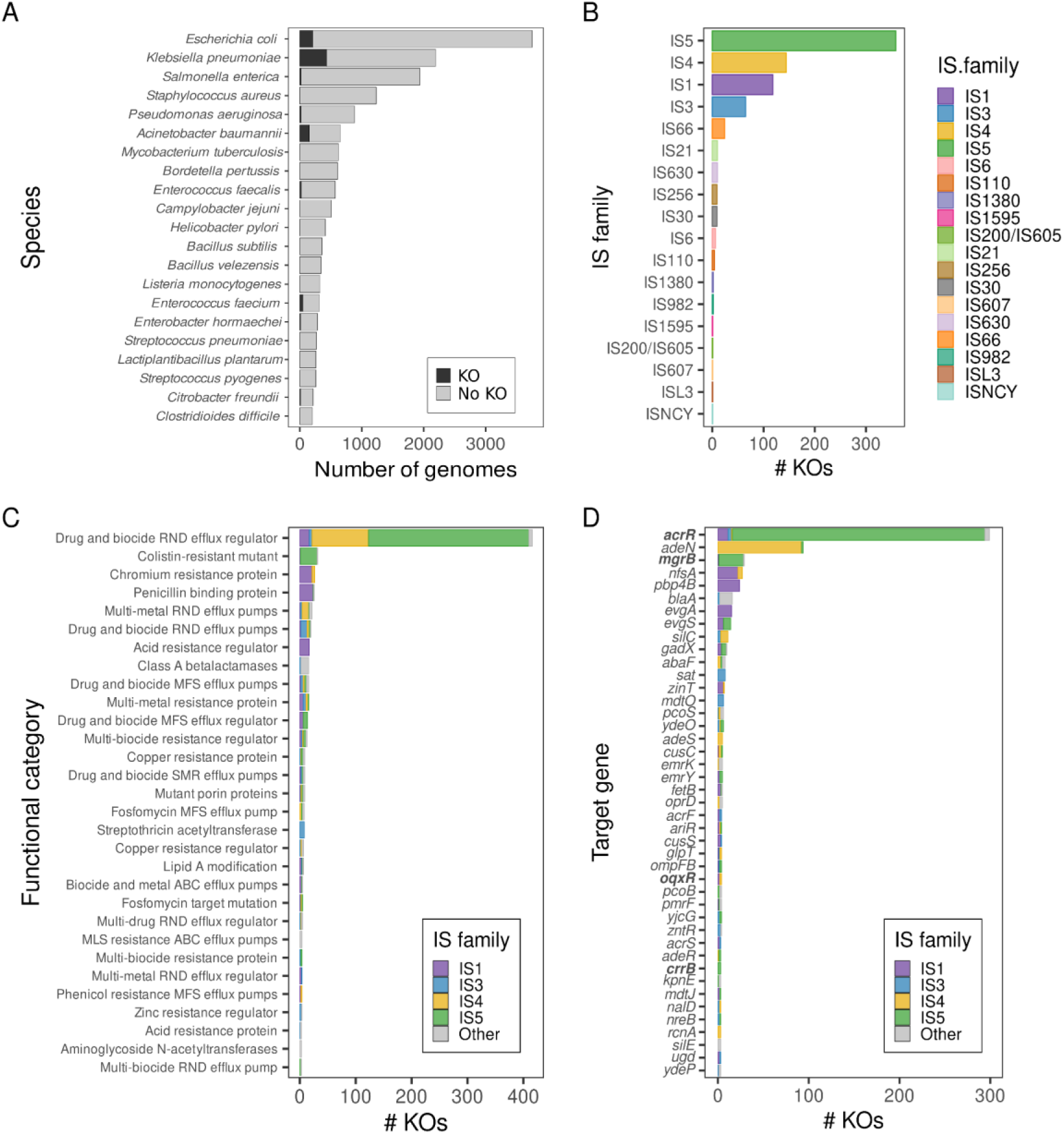
Distribution of IS-mediated KOs in the BV-BRC database. **A)** Top 20 species analyzed from the BV-BRC database. Darker bars indicate those genomes which showed KO in genes from the MEGARES database and our experimental targets. IS-mediated disruptions were specially prevalent in *K. pneumoniae*, *E. coli* and *A. baumannii*, which are species of high clinical relevance. **B)** Main IS families detected disrupting chromosomal genes from MEGARES+experimental targets. IS5, IS4, IS1 and IS3 families were the main families driving KO-AMR. **C)** Main functional categories targeted by IS elements in the ∼42K genomes analyzed from BV-BRC. We observed that most of the targets corresponded to RND efflux pump regulators, followed by colistin resistance related genes. **D)** Main gene targets disrupted by IS elements. Most insertions inactivated the efflux pump regulators *acrR* and *adeN*, which disruption confers MDR phenotypes in Enterobacterales and *A. baumannii* respectively. As expected, we also detected disruptions in *mgrB*. Other genes, such as *nfsA*, which inactivation confers resistance to nitrofurantoin, were among the top detected targets. Bold gene names indicate those genes which were also targeted in our experiments. Main IS families are colored as in panel B. Underrepresented IS families are colored in gray to simplify the visualization.

**Supplementary Figure 5.**
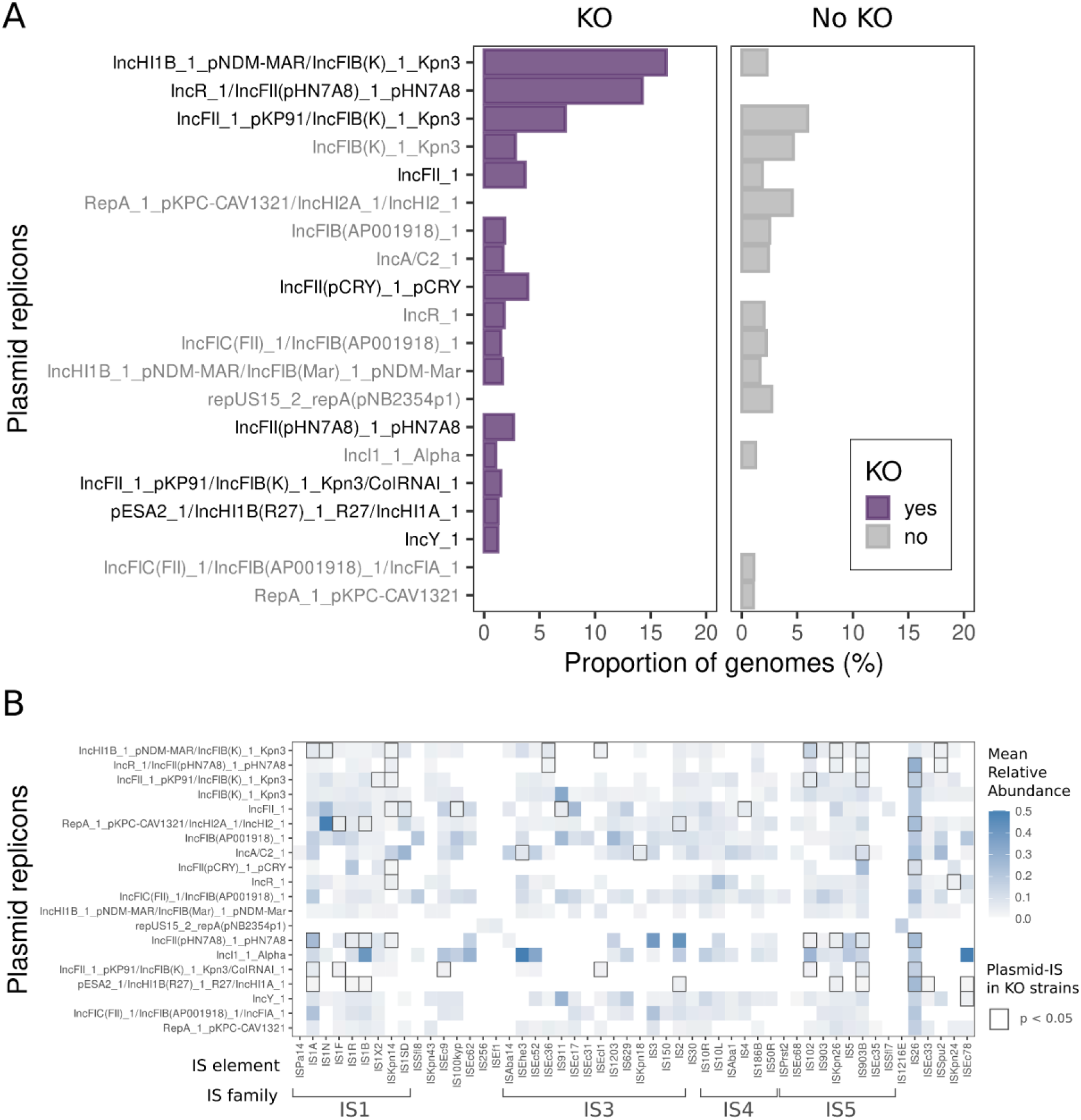
Statistical analyses of plasmid-encoded IS causing KO-AMR. **A)** Frequency of genomes from the BV-BRC encoding a determined plasmid replicon or fusion of replicons in strains with KO-AMR genes inactivated by IS elements (left) or not (right). The replicon names indicated with darker names were found significantly over enriched in strains showing KO-AMR inactivations by ISs (left panel). **B)** Heatmap showing the abundance of plasmid-encoded IS elements. Y-axis indicates the most prevalent plasmid replicons. X-axis shows the IS elements driving the inactivation of the screened KO-AMR genes, sorted by their IS family in alphanumeric order. Only the top families causing KOs are highlighted. Each cell indicates the mean relative abundance (normalized per strain genome) of a given IS element encoded in a plasmid (i.e. if the mean relative abundance value of a Y-plasmid + X-IS combination is 0.1, it indicates that, if present, Y-plasmid encodes, by average, 10% of the X-IS element in a genome). Empty cells correspond to non-detected associations. Highlighted cells with darker stroke lines correspond to plasmid-IS combinations significantly over represented in the subset of strains showing IS-mediated KOs of KO-AMR genes.

**Supplementary Figure 6.**
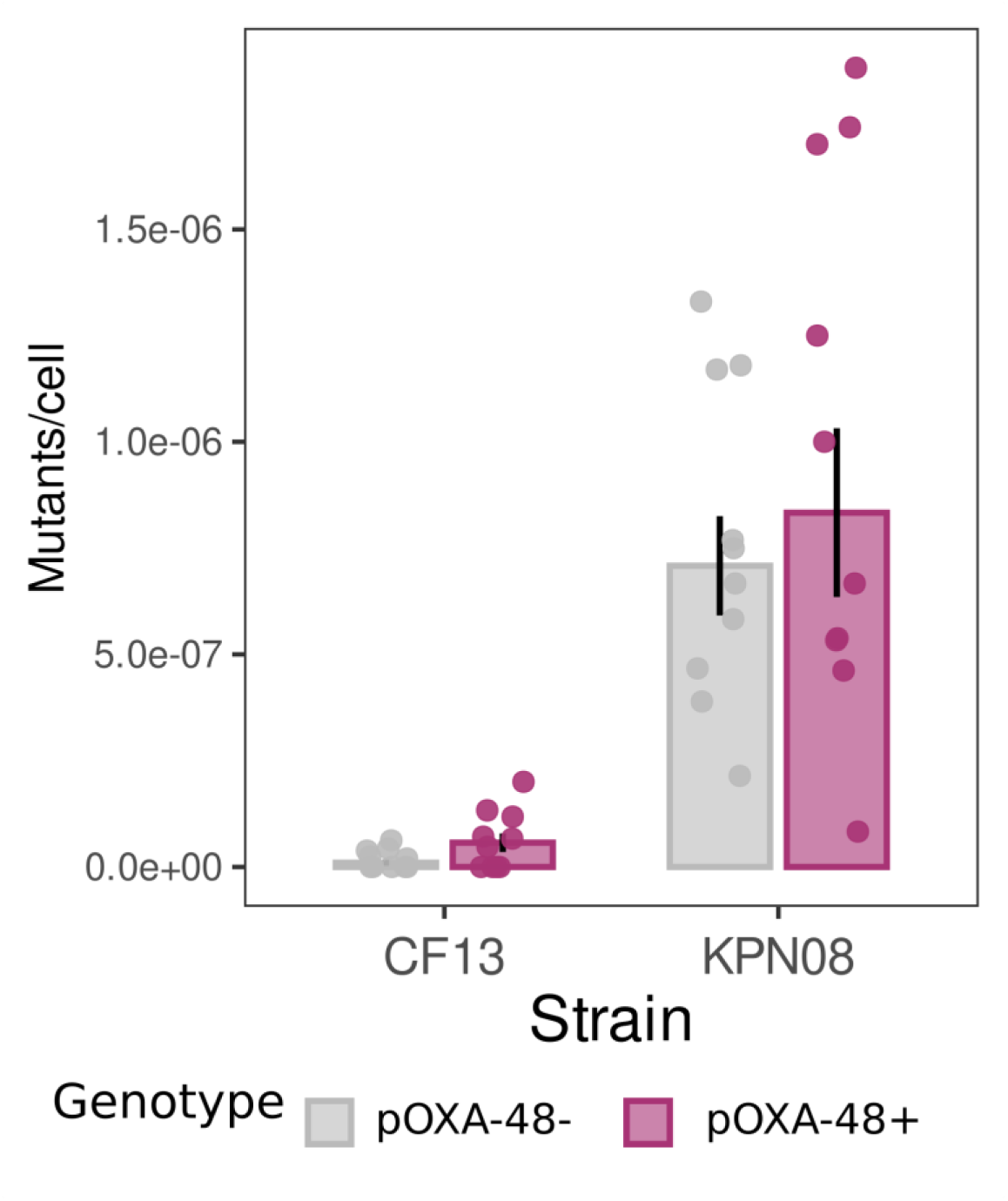
Frequency of colistin resistant mutants in CF13 and KPN08. Frequency of colistin resistance (mutants per cell) for the Enterobacterales members of the community in which pOXA-48 conjugated (*C. freundii* CF13, and *K. pneumoniae* KPN08). Grey bars indicate the frequency of mutants in pOXA-48-free cells, and purple bars in pOXA-48 carrying cells. Dots indicate the individual replicates (n = 10) whereas the bar represents the mean value for each strain genotype. No significant differences were found for any of the strains of the population (t-test, n = 10, p > 0.05 after Bonferroni adjustment).

**Supplementary Figure 7.**
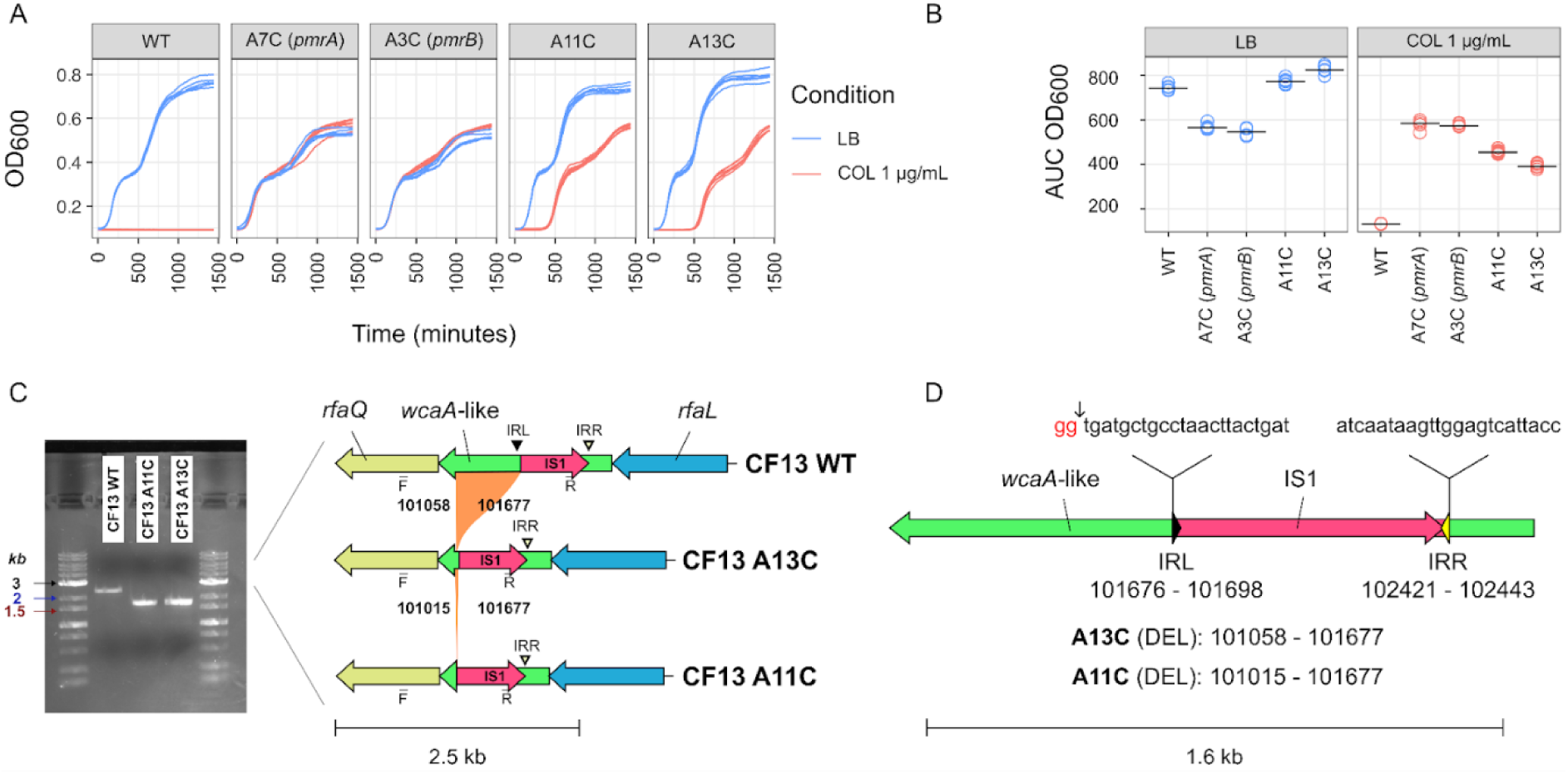
Characterization of colistin-resistant pOXA-48-carrying *Citrobacter freundii* clones isolated from the community experiment. Comparison of wild type C. freundii CF13 (CF13 WT), two of the pOXA-48-carrying COL resistant mutants with mutations in pmrAB (A7C and A3C), and two of the pOXA-48-carrying COL resistant mutants with deletions in the wcaA-like gene putatively mediated by IS1 (A13C and A11C). **A)** Growth curves in liquid media of different *C. freundii* clones. The resistant clones were originally isolated on colistin 1 µg/mL LB agar plates after pOXA-48 invasion (see panels). Vertical axis shows the OD600 and horizontal axis the time in minutes. Each clone was grown 6 times per condition (n=6, blue for LB and red for LB supplemented with colistin 1 µg/mL). **B)** Growth was measured as the area under the growth curve (AUC, vertical axis) for each clone (horizontal axis) after 24 hours. AUC values were calculated from panel A data. Individual dots indicate the AUC value of each replicate and horizontal lines indicate the median value of all replicates (n=6 per clone per treatment). Panels indicate different growth conditions: LB medium (left panel with blue dots) and LB supplemented with colistin 1 µg/mL (right panel with red dots). **C)** Genomic characterization of *wcaA*-like mutations. On the left panel, an agarose gel (1%) electrophoresis shows PCR amplicon fragments from different *C. freundii* clones. Each band corresponds to: CF13 WT (2278 bp), A13C (1660 bp amplicon; 619 bp deletion) and A11C (1616 bp; 662 bp deletion) clones. Note that no mutations in the *wcaA*-like gene were identified in clones A7C and A3C, which carried mutations affecting the *pmrA* and *pmrB* genes, respectively (**S. Table 3**). On the right panel, a DNA sequence comparison of the *wcaA*-like gene region in the three *C. freundii* clones is shown. The deletions detected by genome sequencing were confirmed by PCR and Sanger sequencing. The primers used to amplify the *wcaA*-like gene are indicated by F and R in the lower part of the representation (**S. Table 6**). Gene reading frames are shown as arrows, with the direction of transcription indicated by the arrowheads. Gene names are labeled above the arrows. Arrow colours represent different genes. The orange-shaded areas indicate deletions, with the position (in bp) of each deletion shown relative to the reference sequence (CF13 WT). Scale is in kilobase pairs (kb). **D)** Schematic representation of the *wcaA*-like gene (green) and IS1 element (pink) in CF13 WT. The positions (in bp) and sequences of the IRL (black triangle) and IRR (yellow triangle) are shown in the upper part of the diagram. The lower part illustrates the position of the deletion (in bp and relative to CF13 WT) in the A13C and A11C clones. Note that the IRL sequence is affected by the deletion (2 bp in red).

**Supplementary Figure 8.**
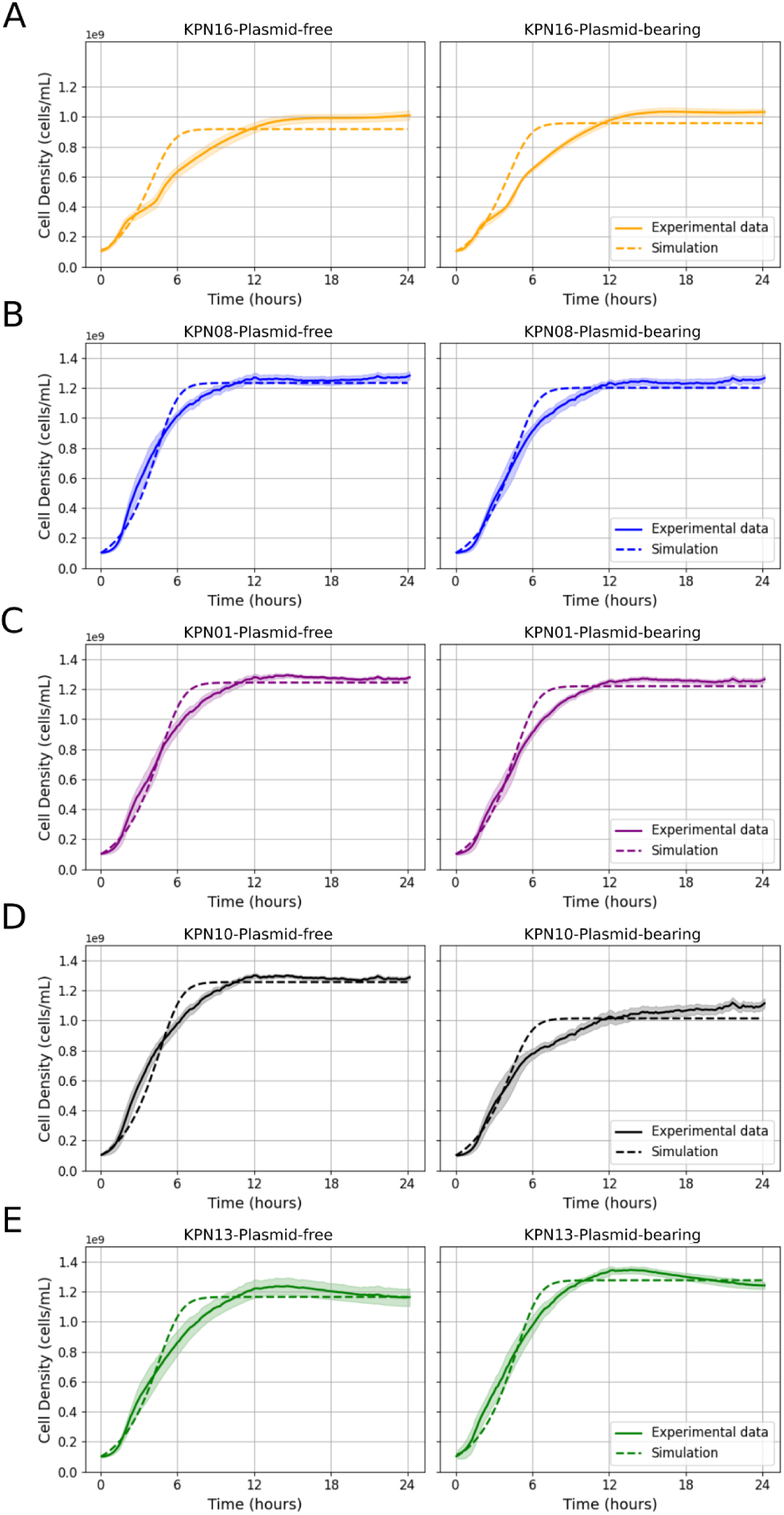
Model parameterization. Comparison between simulations of the stochastic model (dotted lines) and experimental growth data (solid lines) for *Klebsiella pneumoniae* strains grown in isolation. Simulations were performed using best-fit Monod parameters estimated independently for each strain. The left column shows plasmid-free populations, and the right column shows plasmid-bearing populations. Panels correspond to different strains: **(A)** KPN16 (orange), **(B)** KPN08 (blue), **(C)** KPN01 (purple), **(D)** KPN10 (black), and **(E)** KPN13 (green).

**Supplementary Figure 9.**
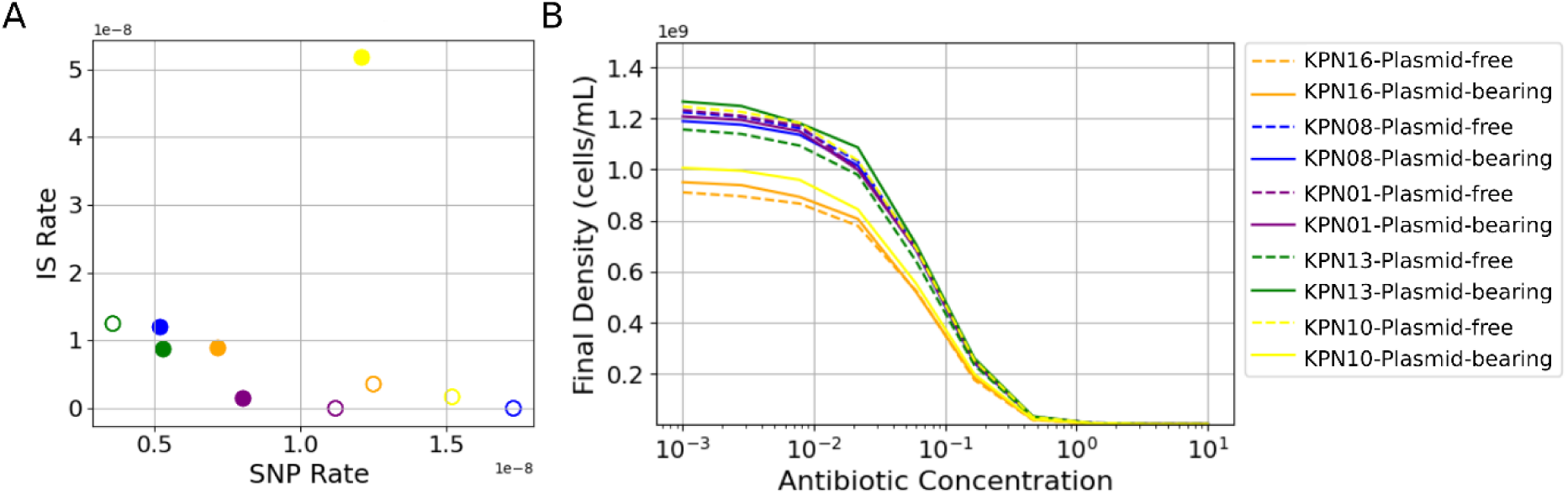
Parameter estimation for mutation, transposition, and antibiotic susceptibility. **A)** Experimentally determined mutation rates (SNPs) and transposition rates (ISs) for each *Klebsiella pneumoniae* strain. Empty circles represent plasmid-free populations; filled circles represent plasmid-bearing populations. **B)** Simulated dose–response curves for plasmid-free and plasmid-bearing populations of K253 KPN16 (orange), KPN08 (blue), KPN01 (purple), KPN10 (yellow), and KPN13 (green) grown in isolation.

**Supplementary Figure 10.**
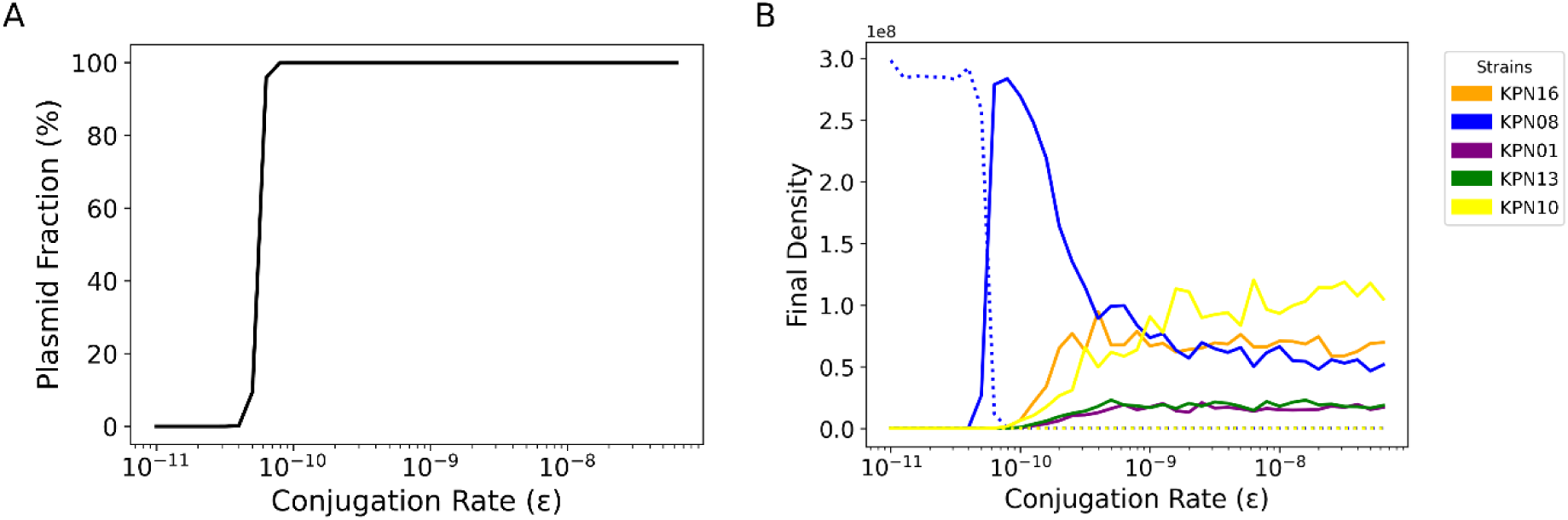
Effect of conjugation rate on plasmid dynamics and community composition. **A)** Relationship between conjugation rate (ε) and plasmid prevalence (%). Increasing ε results in a sharp transition from low to high plasmid frequencies in the population. **B)** Final population densities of different strains after 45 days of evolution under a range of conjugation rates. Strains are represented as KPN16 (orange), KPN08 (blue), KPN01 (purple), KPN10 (yellow), and KPN13 (green). Simulations begin with a community of plasmid-free cells and a small fraction of plasmid-bearing KPN13 cells. As conjugation increases, plasmid-free KPN08 transitions to a plasmid-bearing state, and at high conjugation rates, coexistence of multiple plasmid-bearing strains is observed.

**Supplementary Figure 11.**
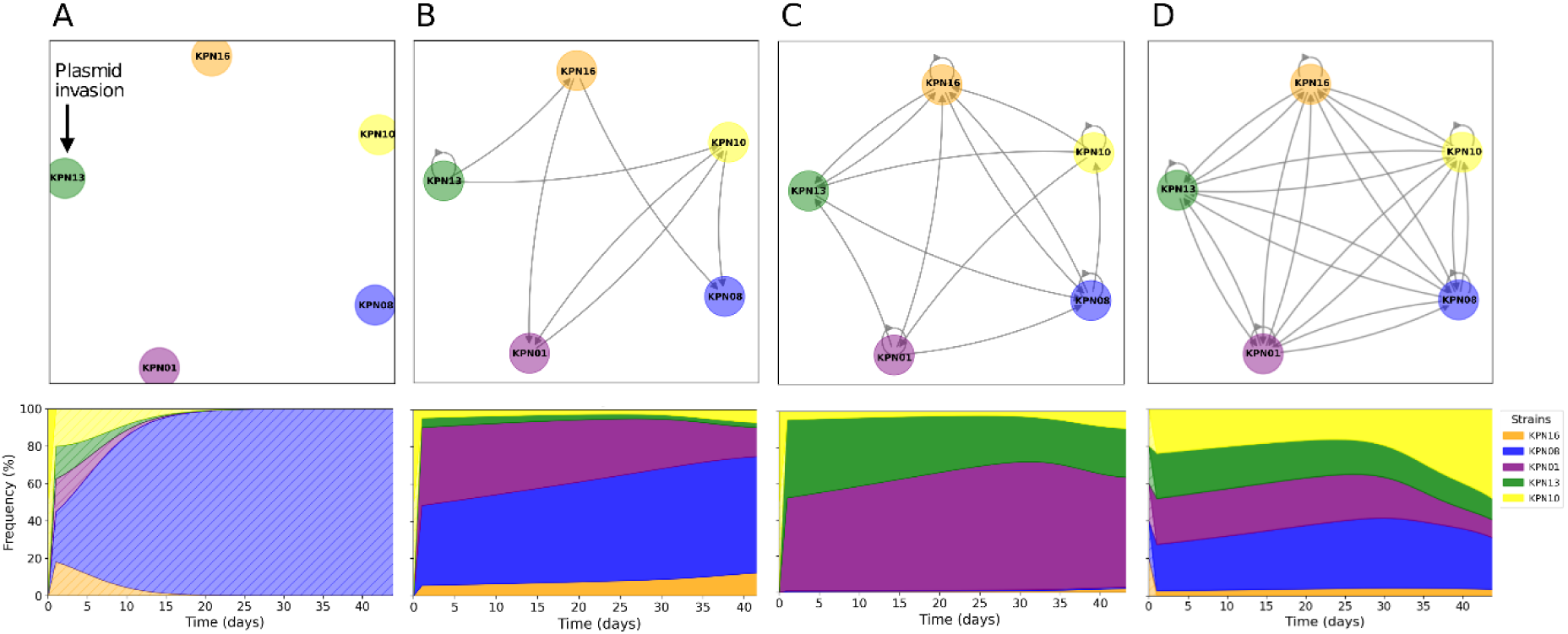
Impact of network structure on plasmid dynamics and community composition. Top row: Examples of plasmid transmission networks with increasing network densities: **A)** 0, **B)** 0.4, **C)** 0.8, and **D)** 1.2. Bottom row: Corresponding Muller plots showing the temporal dynamics of strain frequencies over 45 days. Strains are color-coded as KPN16 (orange), KPN08 (blue), KPN01 (purple), KPN10 (yellow), and KPN13 (green). Simulations begin with a community composed entirely of plasmid-free cells and a small fraction of invading plasmid-bearing KPN13 cells. As network density increases, the population structure changes, promoting plasmid spread and coexistence among multiple strains.

