## Extended Methods for "Plasmids promote antimicrobial resistance through Insertion Sequence-mediated gene inactivation"

**Extended Methods for the manuscript:**

^1^ Centro Nacional de Biotecnología, Centro Nacional de Investigaciones Cientificas (CSIC), Madrid, Spain.

^2^ Center for Genomic Sciences, Universidad Nacional Autónoma de México, 62210 Cuernavaca, México.

^3^ Departamento de Biología Molecular, Facultad de Biología, Universidad Autónoma de Madrid, Madrid, Spain.

^4^ Consorcio de Investigación Biomédica en Red de Epidemiología y Salud Pública (CIBERESP), Madrid, Spain.

* These authors contributed equally and should be considered as first authors.

***Bacterial strains***

All the clinical strains included in this work belong to the R-GNOSIS collection, obtained as part of a surveillance screening program for detecting extended-spectrum ß-lactamases or carbapenemase-producing Enterobacterales in hospitalized patients at the Hospital Universitario Ramón y Cajal (Madrid, 2014-2016), approved by the Hospital Ethics Committee (ref. no, 251/13). The strains selected in the initial screen include 20 *K. pneumoniae* (ST15, ST377, ST432 and ST1427), one *E. coli* (ST131) and one *C. freundii* (ST22). For the community assay, we also used a counterselectable diaminopimelic acid auxotrophic laboratory strain, the *E. coli* 𝛽-3914^1^. Complete description of the strains can be found in **S. Table 1**.

***Antibiotic susceptibility testing***

To select the antibiotics to use in the different experimental assays, we first tested the sensitivity of the selected bacterial strains to 12 antibiotics (**S. Table 1**), including azithromycin, gentamicin, erythromycin, colistin, sulphonamide, fosfomycin, streptomycin, rifampicin, kanamycin, chloramphenicol, tetracycline and ciprofloxacin. To do so, we performed antibiograms in Luria Bertani (LB) agar plates (Condalab). First, we individually inoculated each strain directly from the stock in 2 mL of LB. Then, we incubated the cultures for 18 hours at 37ºC and constant shaking (Synergy HTX Multi-Mode Reader; BioTek Instruments; 200 rpm). After the incubation, we plated each strain on LB agar and placed the antimicrobial susceptibility disks (Bio-Rad) using a 16 disk dispenser for cartridges (Bio-Rad) for all the antibiotics tested. We incubated the samples for 18 hours at 37ºC. Finally, we measured the inhibition halo diameter and counted the number of spontaneous mutants inside it (**S. Table 1**).

**Plasmid stability and antagonistic interactions.** We discarded plasmid loss for each plasmid/community member by streaking from stocks the LB plates supplemented with the corresponding antibiotics and incubating them overnight at 37ºC. We picked three independent colonies per genotype to start 5 mL LB cultures. Next, after an overnight incubation at 37°C with shaking at 250 rpm, we performed serial dilutions (from 1:10 to 1:10^7^) in 0.9% NaCl and plated on LB agar plates with and without antibiotics. We incubated the plates at 37°C to estimate CFUs for each genotype and for each media. In parallel, we transferred 5 µL of the overnight cultures into 50 mL tubes containing 5 mL of fresh LB media. We incubated these cultures under the same conditions as before, serially diluted in 0.9% NaCl and plated on LB agar plates with and without antibiotics. After overnight incubation at 37°C, we estimated CFUs again. We discarded possible antagonistic interactions between members of the community by growing them together on LB agar plates. Briefly, we streaked each member of the community in contact on an LB agar plate. We incubated the plate for 24 and 48 hours to discard possible interactions.

**Variant calling.** To detect mutational events in resistant mutants, we performed the variant calling of the Illumina short reads using breseq v.39.0.^6^ against the assembled reference genome of each strain. For the population WGS data obtained from the high-throughput and community assays, we used the ‘-p’ flag to detect polymorphic mutations. To analyze the results, we developed a Python script (Python v.3.8) to parse the HTML results given by breseq to obtain xlsx files (see ‘Code availability’) and simultaneously compare mutations present in the samples by merging their reports into an individual table per strain. In all experiments, we considered all mutational events that could confer a resistant phenotype. For population samples from the high-throughput and community assays, we considered a mutation to be fixed in a population when its frequency reached 50%. We discarded mutations with a frequency below 10% and only considered those above 20% frequency to contribute to resistance. To visualize and interpret the deletion in *C. freundii* samples, we used IGV v.2.19.

***Computational model***

**Stochastic Model of Plasmid Dynamics and Mutation Accumulation.** We developed a stochastic, multi-species model based on the Gillespie algorithm to simulate the evolutionary dynamics of bacterial populations under selection. The model captures how cellular processes (bacterial growth and death), plasmid dynamics (segregation and conjugative transfer), and evolutionary events (mutation and transposition) are influenced by environmental and physiological factors. We modeled all events as discrete stochastic transitions, with propensities determined by strain-specific parameters, including growth rates, mutation and transposition rates, antibiotic susceptibility, and conjugation efficiency, as well as population size ($N$), resource availability ($R$), and antibiotic concentration ($A$). Growth was modeled using monod kinetics under a homogeneous environment, with resources equally accessible to all cells. We incorporated antibiotic pressure by modulating death rates via a saturating dose–response function. Plasmid segregation occurred stochastically at cell division, resulting in the plasmid's loss and the cell's reclassification into the plasmid-free subpopulation. We modeled conjugation as a mass-action process, with successful transfer dependent on the conjugation rate and the permissiveness of the recipient cell. The simulated community was composed of 5 species, each partitioned into plasmid-free and plasmid-bearing subpopulations. The number of accumulated SNPs further stratified these subpopulations and IS transpositions, allowing the model to track mutation dynamics and clonal interference within each species. Mutation and transposition events occurred independently and irreversibly, incrementing a cell’s mutation or IS count by one unit. We assumed all mutations affect phenotype, either conferring resistance or imposing a fitness cost. Plasmid transfer between strains is determined by a directed $M \times M$ adjacency matrix ε_ij_ , where each element defines the conjugation rate from strain $i$ to strain $j$. In the base case, we assume a fully connected network in which all strains can transfer plasmids to all others at a uniform rate $\varepsilon>0$. We estimated model parameters independently for each bacterial strain using experimental data obtained from monoculture growth assays and mutation rate measurements. SNP and IS mutation rates were obtained from fluctuation assays, where mutation frequencies were quantified using whole-genome sequencing of replicate cultures. To infer monod kinetics parameters, the intrinsic growth rate (*μₘₐₓ*) and the half-saturation constant (*Kₛ*), we fitted a simplified, single-strain version of the model that excluded mutation, segregation, and conjugation processes. We performed parameter optimization using the L-BFGS-B algorithm to minimize the discrepancy between experimental and simulated growth curves, quantified by the difference in final optical density (OD). We conducted simulations under initial resource concentrations and time frames consistent with the experimental conditions (T = 24 h), and we enforced biologically plausible parameter bounds during optimization. We inferred strain-specific death rate parameters under antibiotic pressure by fitting model simulations to experimental IC₉₀ values, defined as the concentration at which 90% of the population is inhibited. We simulated growth and survival across a range of antibiotic concentrations and optimized the drug-dependent mortality function to match the observed susceptibility profiles.

**Simulating Evolutionary Dynamics in Serial Transfer Experiments.** We ran simulations in discrete 24-hour cycles, representing daily growth phases. At the end of each cycle, populations were diluted by a fixed factor (1:100) to simulate serial transfer conditions, maintaining relative strain and plasmid frequencies while reducing absolute cell numbers. Resource and antibiotic concentrations were replenished to their initial values at each transfer, allowing for consistent environmental conditions across days. This setup allowed us to track plasmid dynamics and the accumulation of mutations over 45 simulated days. We performed two classes of *in silico* experiments. In the first, we fixed the network topology as a complete graph in which all strains could exchange plasmids with one another and we varied the conjugation rate (ε). This allowed us to identify thresholds in conjugation efficiency associated with changes in plasmid prevalence and mutation spectra. In the second set of experiments, we fixed ε=10⁻^9^. we represented each resulting configuration as a directed transmission matrix ε_i_, encoding the conjugation rate from strain $i$ to strain $j$. We randomly removed edges from the complete graph to systematically vary network topology while preserving global connectivity. Network density was defined as the fraction of possible directed edges present between distinct strains. To quantify evolutionary outcomes, we tracked the population dynamics of each strain over time, including total cell counts, plasmid status, and the accumulation of SNPs and ISs in both plasmid-free and plasmid-bearing subpopulations. Mutational events were irreversible and strain-specific, and were recorded throughout the simulations.
